# Investigating the effect of changes in model parameters on optimal control policies, time to absorption, and mixing times

**DOI:** 10.1101/2023.01.23.525286

**Authors:** Kathleen Johnson, Daniel Plaugher, David Murrugarra

## Abstract

Many processes in biology and medicine have been modeled using Markov decision processes which provides a rich algorithmic theory for model analysis and optimal control. An optimal control problem for stochastic discrete systems consists of deriving a control policy that dictates how the system will move from one state to another such that the probability of reaching a desired state is maximized. In this paper, we focus on the class of Markov decision processes that is obtained by considering stochastic Boolean networks equipped with control actions. Here, we study the effect of changes in model parameters on optimal control policies. Specifically, we conducted a sensitivity analysis on optimal control policies for a Boolean model of the T-cell large granular lymphocyte (*T-LGL*). For this model, we quantified how the choice of propensity parameters impacts the effectiveness of the optimal policy and then we provide thresholds at which the effectiveness is guaranteed. We also examined the effect on the optimal control policies of the level of noise that is usually added for simulations. Finally, we studied the effect on changes in the propensity parameters on the time to absorption and the mixing time for a Boolean model of the Repressilator.

## 1 Introduction

Boolean network (BN) models are becoming increasingly popular and have been successfully used to model important biological systems such as the bistable behavior of the *lac operon* [1], the yeast-to-hyphal transition of the yeast *Candida albicans* [2], the interaction of pancreatic cancer cells with their microenvironment [3, 4], and they have been proposed as appropriate models for understanding the evolution of cellular differentiation [5]. Furthermore, there is a growing collection of BN models in the *Cell Collective* database [6] covering processes such as signaling, regulation, and cancer.

To account for the stochasticity in various molecular processes, several versions of stochastic Boolean networks exist including Probabilistic Boolean networks (PBN) [7, 8], perturbed Boolean networks (BNp) [9, 10], and Probabilistic Edge Weights operators (PEW) [11]. This paper will focus on the stochastic framework that was introduced in [12] referred here as Stochastic Discrete Dynamical Systems (SDDS). The stochasticity usually originates from the updating schedule or from an added perturbation to the functions. Standard updating schedules include synchronous update where all nodes are updated at the same time, and asynchronous update, where a random set of nodes are updated at each time step. Consequently, a synchronous update produces deterministic dynamics, while an asynchronous update generates stochastic dynamics. The SDDS framework introduces stochasticity by assigning two propensity parameters to each function in the Boolean network. The PBN framework considers multiple functions for each node where one function is selected at each step using a probability distribution.

A Markov decision process (MDP) is a mathematical modeling framework for decision making [13, 14]. An optimal control policy for an MDP provides an action for each state of the system such that a certain optimality criterion is achieved. Many methods exist for obtaining optimal control policies such as the infinite-horizon method with a discount factor [15, 16]. The optimal control policies are usually obtained using the value iteration method for approximating the solution of Bellman’s equation [17, 18].

The main contributions of this paper will be presented in the Results section, which is divided in two parts. In the first part, we study the effects of changes in the propensity parameters of the SDDS framework on the optimal control policies for a Boolean model of the T-cell large granular lymphocyte (*T-LGL*) [19]. For this model, we show how the choice of propensity parameters impacts the effectiveness of the optimal policy and then we provide thresholds for which the effectiveness is guaranteed. We also examined the effect of the level of noise that is usually added for simulations on the optimal control policies. In the second part of the Results section, we discuss the effects of the propensities on the long-term dynamics of a Boolean model of the Repressilator. For this model, we studied the effect of changes in the propensity parameters on the time to absorption and the mixing time.

This paper is structured in the following way. In Section 2, we introduce the modeling framework SDDS and their associated Markov decision processes. In Section 3.1 we present the results of the effects of changes in the propensities on control policies for the T-LGL model. In Section 3.2, we discuss the long term dynamics effects of propensities on a Boolean model of the Repressilator. In Section 4 we discuss our main findings and provide some conclusions.

## 2 Methods

In this section we describe the stochastic modeling framework SDDS and the optimal control method that we will use in this paper.

### 2.1 Stochastic Discrete Dynamical Systems

Consider *n* variables *x*_1_, …, *x*_*n*_ that can take values in finite sets *X*_1_, …, *X*_*n*_. Let *X* = *X*_1_ *×· · · × X*_*n*_ be the Cartesian product. An SDDS in the variables *x*_1_, …, *x*_*n*_ is defined as a collection of *n* triples

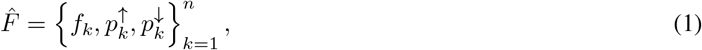

where for *k* = 1, …, *n*,

- *f*_*k*_ : {0, 1}^*n*^ → {0, 1} is the update function for *x*_*k*_,
- 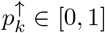 is the activation propensity,
- 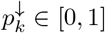 is the degradation propensity.

The activation parameter 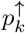 represents the probability that an activation of *x*_*k*_ occurs, given that *f*_*k*_ would activate *x*_*k*_. That is, if *x*_*k*_(*t*) = 0, and *f*_*k*_(*x*_1_(*t*), …, *x*_*n*_(*t*)) = 1 then *x*_*k*_(*t* + 1) = 1 with probability 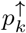. The degradation propensity 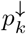 is defined similarly.

We can represent an SDDS as a Markov chain with the transition probability from *x* = (*x*_1_, …, *x*_*n*_) to *y* = (*y*_1_, …, *y*_*n*_) is given by

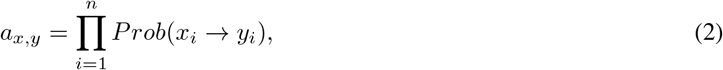

where *Prob*(*x*_*i*_ *→ f*_*i*_(*x*)) is the probability that *x*_*i*_ will change its value and is given by

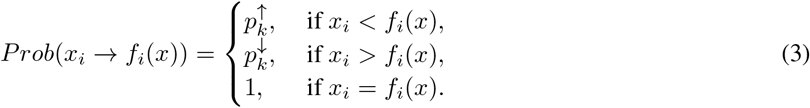

The probability that *x*_*i*_ will maintain its current value is given by

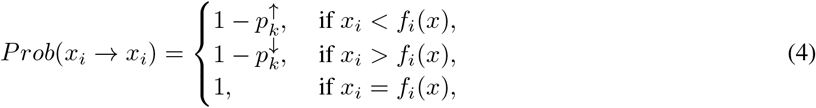

for all *i* = 1, …, *n*. Note that *Prob*(*x*_*i*_ *→ y*_*i*_) = 0 for all *y*_*i*_ *∉ {x*_*i*_, *f*_*i*_(*x*)*}*. The transition matrix is

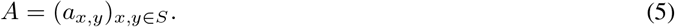

### 2.2 Boolean Networks

Boolean networks are discrete dynamical systems. Specifically, consider *n* variables *x*_1_, …, *x*_*n*_ each of which can take values in {0, 1}. Then, a synchronously updated Boolean network is a function *F* : (*f*_1_, …, *f*_*n*_) : {0, 1}^*n*^ *→* {0, 1}^*n*^, where each coordinate function is a discrete function on a subset of {*x*_1_, …, *x*_*n*_}, which represents how the future value of the *i*th variable depends on the present values of the variables. We note that setting all propensity parameters 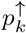 and 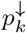 to 1.0 in the SDDS framework will result in a synchronous Boolean network.

### 2.3 Markov Chains

A discrete-time Markov chain is a sequence of random variables *χ*_1_, *···, χ*_*n*_, *···* with the Markov property: *Prob*(*χ*_*n*+1_ = *x*| *χ*_1_ = *x*_1_, *···, χ*_*n*_ = *x*_*n*_) = *Prob*(*χ*_*n*+1_ = *x*| *χ*_*n*_ = *x*_*n*_). That is, the probability of moving to the next state depends only on the current state and not on the previous states. We note that Boolean networks are Markov chains.

### 2.4 Markov Decision Processes

Here we will describe a Markov Decision Process associated to the SDDS framework [18]. To use tools from Markov Decision Processes with SDDS, we consider the underlying Boolean network *F* and corresponding wiring diagram *G* associated with the SDDS 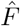. The wiring diagram *G* has nodes *x*_1_, …, *x*_*n*_, and there is a directed edge from *x*_*i*_ to *x*_*j*_ if *f*_*j*_ depends on *x*_*i*_. The edge and node controls can be encoded in the Boolean network 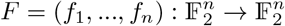.

Let *U* be the set of all possible control inputs, where 0 represents no control. We encode the edge and node controls using the function 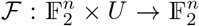 such that *ℱ* (*x*, 0) = *F* (*x*) returns the original Boolean network. To include control in an SDDS, we replace the functions of *f*_*k*_ with *ℱ*_*k*_ : *{*0, 1*} × U → {*0, 1*}*. With edge control, *u*_*i,j*_ refers to control of the edge *x*_*i*_ *→ x*_*j*_. For node control, *u*_*i*_ refers to node *x*_*i*_. In both edge and node control, setting *u* to zero represents an inactive control and setting *u* to one represents removal of the corresponding edge or node.

A control action *a* is a combination of control edges and nodes being applied to the Boolean network simultaneously at a given time step. A set *E* of control edges and set *V* of control nodes can be used to define *a* as an array of binary elements of size |*U* | = |*E*| + |*V* |. Let *A* = *{*0, 1*}*^|*U* |^ be the set of all possible actions.

Now, we define an MDP for the SDDS 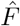 using these control actions. A Markov decision process for SDDS has four components: the set of states *S*, the set of control actions *A*, the transition probability 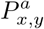 associated with action *a* at state *x* that will lead to state *y*, and the associated costs *C*(*x, a, y*).

The new SDDS resulting from the application of action *a* is 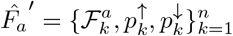. The transition probability 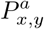 from state *x* to *y* associated with action *a* is computed with equation 2 where *f*_*k*_ is replaced by 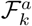. The cost function *C*(*x, a, y*) describes the penalties for going from state *x* to *y* using action *a*. Thus the cost is given as the sum of the cost for the action and one for the state:

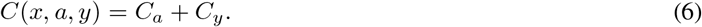

The action cost *C*_*a*_ = *c*_*v*_*N*_*v*_ + *c*_*e*_*N*_*e*_ where *c*_*v*_ and *c*_*e*_ are the penalties for the control nodes and edges respectively, and *N*_*v*_ and *N*_*e*_ are the number of applied control nodes and edges in the given action *a*. The state cost

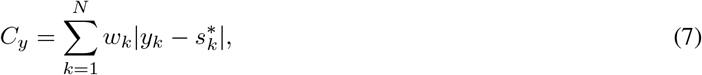

where *s*^*\**^ is the desired state and *w*_*k*_ are the weights of each edge.

### 2.5 Optimal Control for SDDS

This section reviews a method for optimal control for Markov decision processes called the infinite-horizon method with a discounting factor as presented in [18].

A deterministic control policy *π* is a set *π* = {*π*_0_, *π*_1_, …} where each *π*_*t*_ associates a state *x*(*t*) to an action *a* at time step *t*. Given a state *x ∈ S*, a control policy *π* and a discounting factor *γ ∈* (0, 1), the cost function *V* ^*π*^ associated with *π* is defined as

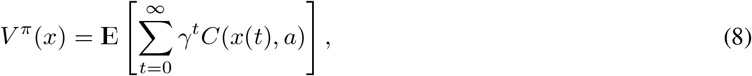

where *C*(*x*(*t*), *a*) represents the expected cost at time step *t* for executing the policy from state *x*. The cost at step *t* is defined by

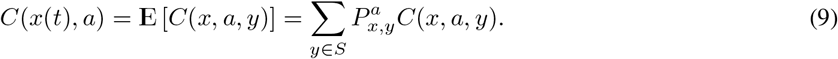

The optimal cost function *V* ^*\**^(*x*) associated with the optimal policy *π*^*\**^ is

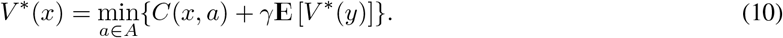

Because this function satisfies Bellman’s equation [13, 14], we can use the value iteration algorithm to calculate the optimal control policy. The equation for the value iteration algorithm is

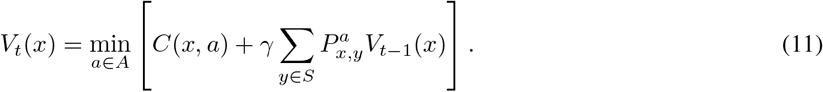

For any initial bounded cost function *V*_0_, the recursion will converge to *V* ^*\**^.

### 2.6 Time to Absorption

Consider the transition matrix of a Markov chain in canonical form,

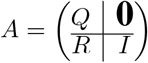

where **0** is the zero block matrix and I is the identity submatrix. The fundamental matrix *N* is defined as the inverse of (*I − Q*). That is, *N* = (*I − Q*)^*−*1^. The time to absorption for a transient state *j* is defined as the expected number of steps before absorption and can be calculated as the sum of the *j*th row of *N* (see Theorem 11.5 of [20]).

### 2.7 Mixing Time

The mixing time *t*_*mix*_ of a Markov chain is the number of steps required for the chain to be within a fixed threshold of its stationary distribution [21]. More precisely, we define

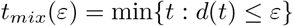

where *d*(*t*) is the total variation distance which is defined by 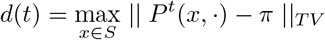. The maximum in the previous formula is taken over all states in the state space *S, P* represents the transition matrix, and *π* denotes the stationary distribution.

Several estimates and bounds exist for mixing times of Markov chains. For irreducible Markov chains, one can show that (see Chapter 12 of [21])

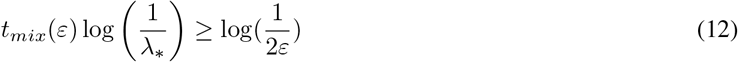

where *λ*_*\**_ is the eigenvalue with the second highest absolute value. In this paper, we will use the following estimate which can be obtained from Equation 12

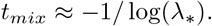

## 3 Results

In this section we provide results from our analysis using a Boolean model where optimal control policies have been calculated. We will apply the optimal control method described in Section 2.5 and study its effectiveness at different propensities and discount factors. We will also study the effect of changes in the level of noise for simulations.

### 3.1 T-LGL model

Here we consider the reduced model of the T-cell large granular lymphocyte (*T-LGL*) [19] with nodes and functions described in Figure 1. This model has two fixed points: 000001 and 110000, where we represented a state of the system by *x*_1_*x*_2_*x*_3_*x*_4_*x*_5_*x*_6_ for each *x*_*i*_ *∈* {0, 1} for *i* = 1,, 6. Here, a state of 0 means that the corresponding gene is inactive while the state 1 indicates that the corresponding gene is active. Thus, the state 000001 represents the desired state (with the apoptosis node ON, *x*_6_ = 1) and 110000 represents the disease state (with the apoptosis node OFF, *x*_6_ = 0). The optimal control problem here is that we want to obtain a policy to ensure the system transitions to the desired state. Note that this system has 64 possible states (since 2^6^ = 64) as indicated in the *x*-axis of Figure 2.

**Figure 1:**
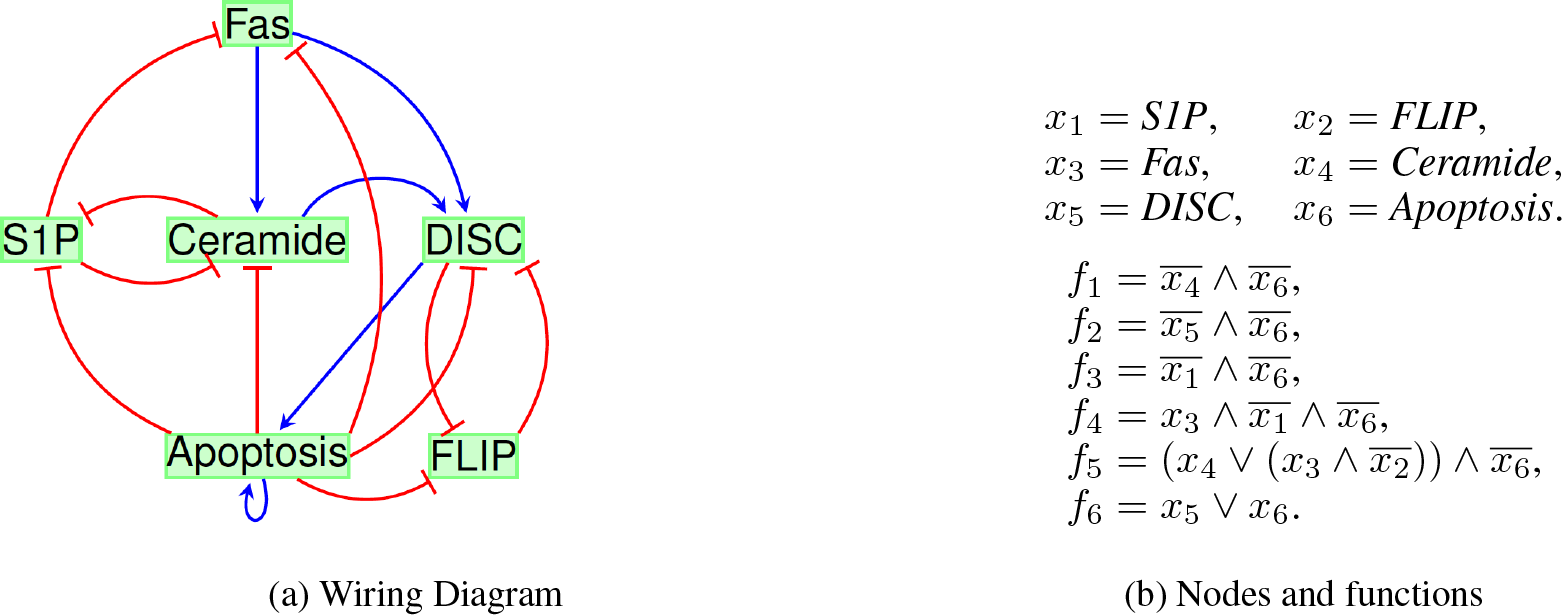
Reduced *T-LGL* network adapted from Saadatpour et al., *PLoS Comput Biol*., 7(11), 2011.

**Figure 2:**
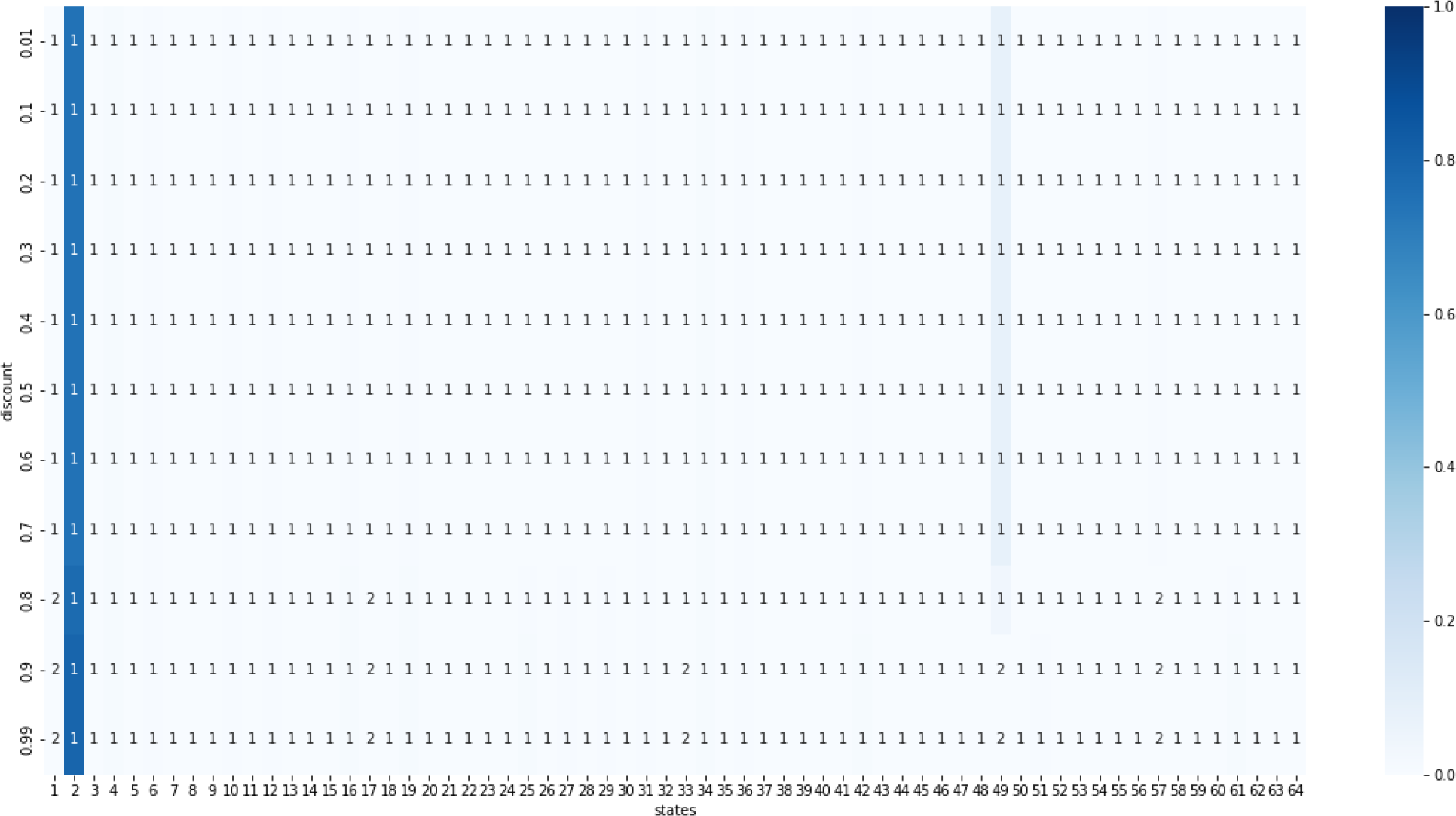
Heatmap of stationary distributions given all propensities are 0.9 using single-step optimal policies from the value iteration method. The states are on the *x*-axis and the *y*-axis gives the discount used to find the policy. At a discount of 0.8 or higher, the stationary distribution consistently goes to state 2 (000001), so this is when the policy becomes effective. The numbers (1 or 2) shown in each row correspond to the action specified by the policy for the corresponding state below the number. The two actions are: (1) no intervention and (2) knock-out of *S*1*P*. The shading gradient on the right of the heatmap correspond to the proportion of time the process ends in one of the possible states.

Given a Boolean network model, different methods for phenotype control can be used to identify edge and node controls including [22, 23, 24]. For the *T-LGL* model discussed in this section, we use the approach in [22] to identify control nodes that can stabilize the system into one of its attractors. Specifically, we will use the control node *S*1*P* = *OFF*. This control was predicted to be 100% effective in [19]. That is, the application of this control (i.e, setting the node *S*1*P* to 0) will result in a controlled network with a single desired fixed point (i.e., 000001). Thus, at each state of the system, we will get access to two actions: (1) no intervention and (2) knock-out of *S*1*P*. Here we say that a policy is effective if this always leads the model to the fixed point 000001. In what follows, we study the effect of changes in the propensity parameters 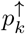 and 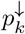 of the SDDS framework on optimal control policies. Two commonly used set of values for the propensities are 0.9 and 0.5. Setting all propensity parameters 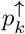 and 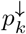 to 0.9 gives equal (90%) chance of update to each variable and therefore this model will highlight the synchronous transitions. Likewise, setting all propensities 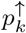 and 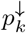 to 0.5 gives equal (50%) chance of update to each variable and therefore this model will be closer to the fully asynchronous case.

In all heatmap figures we consider that state 2 (i.e., 000001) as the desired state while state 49 (i.e., 110000) as the undesired state. The numbers (1 or 2) shown in each row of the heatmap correspond to the action specified by the policy for the corresponding state below the number. The two actions are: (1) no intervention and (2) knock-out of *S*1*P*. The shading gradient on the right of the heatmaps correspond to the proportion of time the process ends in one of the possible states.

In our first result, shown in Figure 2, we have set all propensities 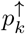 and 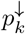 to 0.9. Allowing the discounting factor to vary, we see the policy becomes effective when the discount is at least 0.8. This coincides with a change in the action at the disease node 110000 (state 49 in the horizontal axis) that indicates that the intervention is active.

Setting the propensities 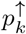 and 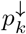 to 0.5 as in Figure 3, we see that the policy is never effective. Note that the action at state 110000 (state 49 in the horizontal axis) does not change to include the control node (i.e., action number 2) for any discounting factor.

**Figure 3:**
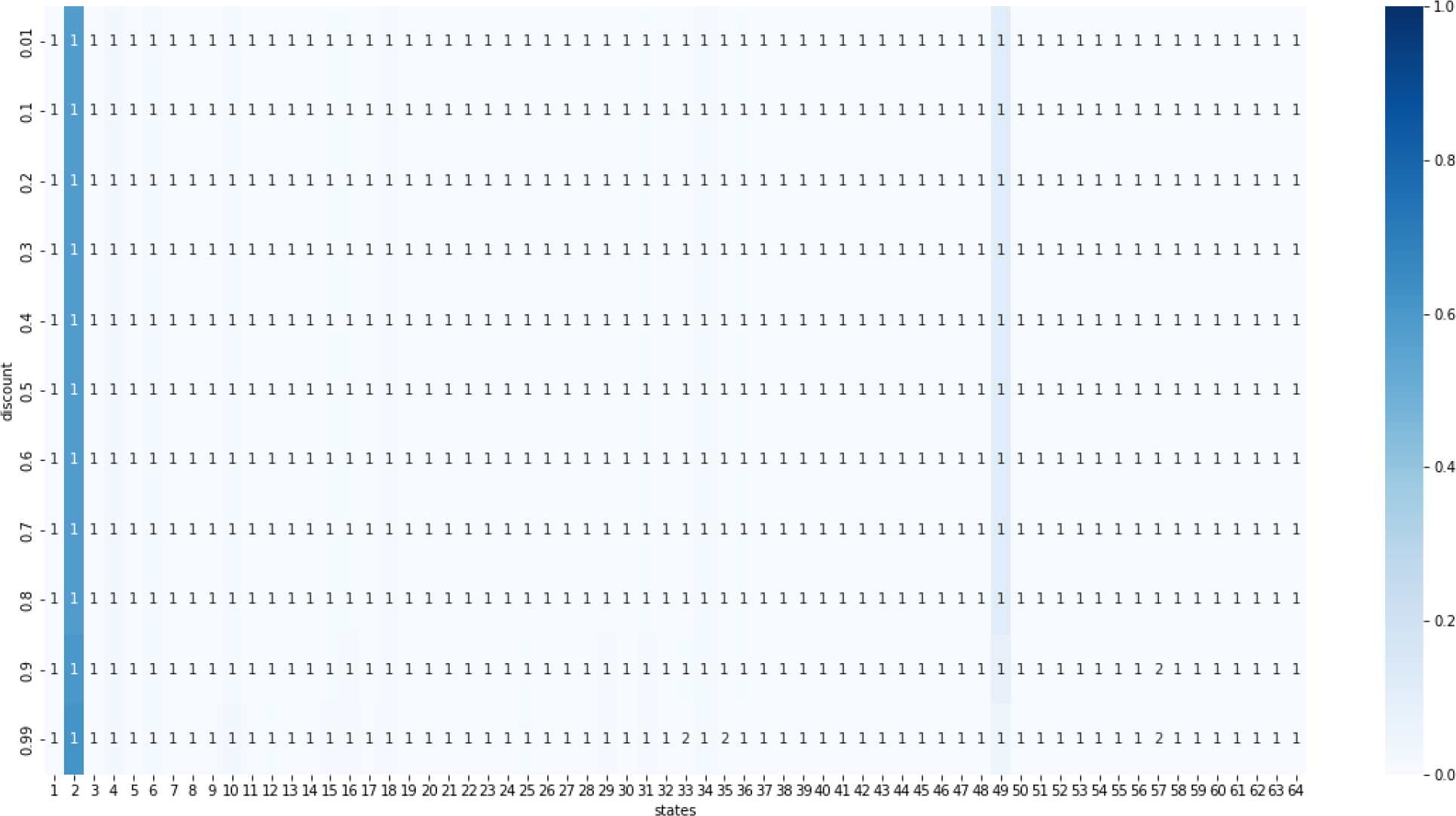
Heatmap of the stationary distribution given all propensities are 0.5 using single-step optimal policies from the value iteration method. The states are on the *x*-axis and the *y*-axis gives the discount used to find the policy. The policies for these choice of parameters are always ineffective.

For our third case, we considered a set of propensity parameters given in Table 5.3 of [18] (we reproduced these propensities here in Table 1). These propensities were obtained from a genetic algorithm to maximize the probability of reaching state 110000. When the value iteration is applied to the model with these propensities as in Figure 4, we see that the policy is never effective. We also note that the action at state 110000 (state 49 in the horizontal axis) does not change to include both interventions for any discounting factor.

**Table 1:** Propensity values computed with a genetic algorithm in [18].

| | $x_1$ | $x_2$ | $x_3$ | $x_4$ | $x_5$ | $x_6$ |
| --- | --- | --- | --- | --- | --- | --- |
| $p_i^\uparrow$ | 0.4792 | 0.4826 | 0.7098 | 0.3911 | 0.0549 | 0.0314 |
| $p_i^\downarrow$ | 0.0274 | 0.9602 | 0.5195 | 0.7758 | 0.8373 | 1.0000 |

**Figure 4:**
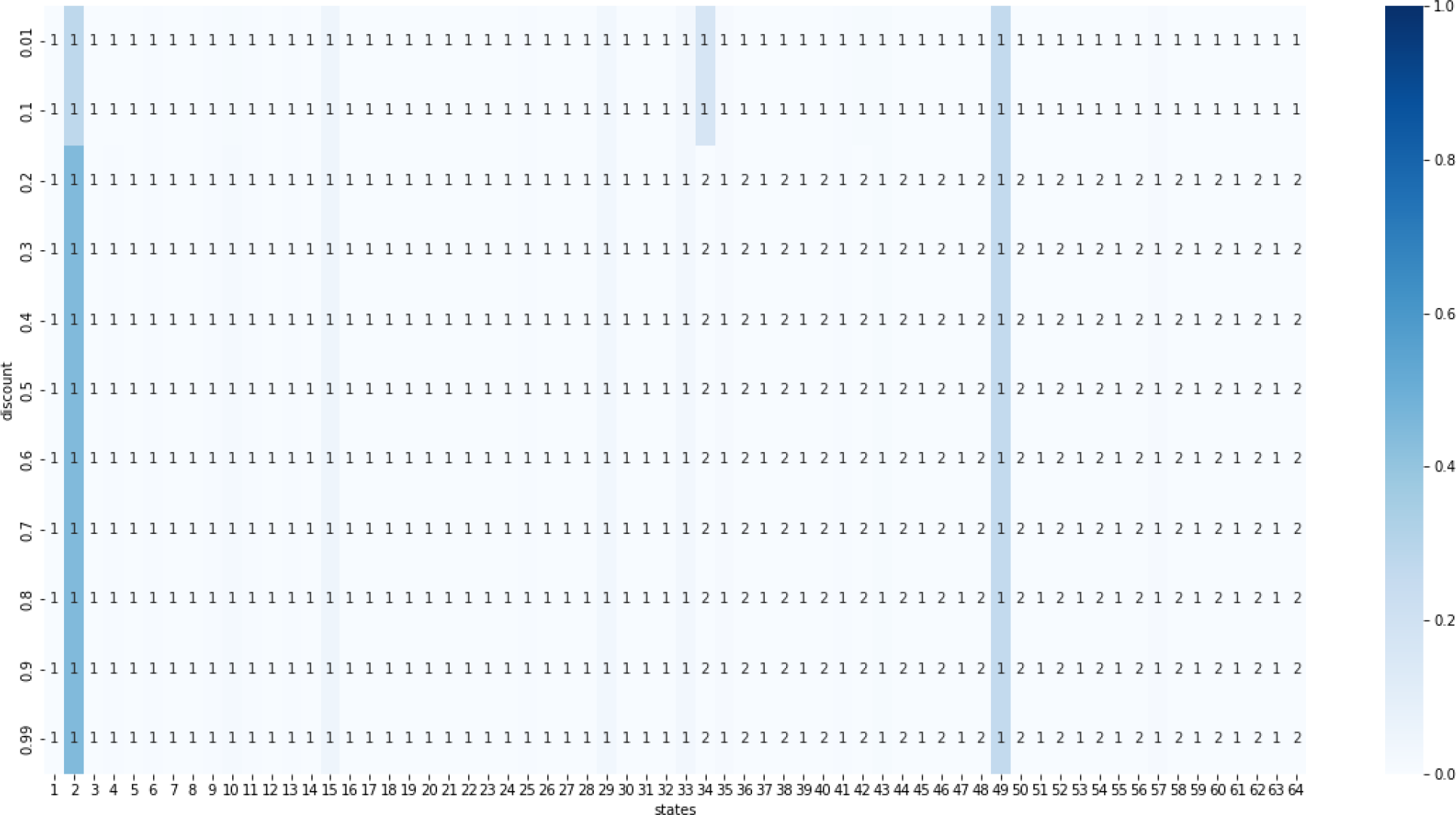
Heatmap of the stationary distribution given propensities from Table 1 using single-step optimal policies from the value iteration method. The states are on the *x*-axis and the *y*-axis gives the discount used to find the policy. The policies for these choice of parameters are always ineffective.

Next, we examined the effect of the noise in the results shown in Figures 2-4. In Figures 5-7 we considered noise levels of 1%, 5%, 10%, and 15% when calculating the optimal policies. In the first case, when we set all propensities 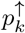 and 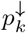 to 0.9, allowing the noise level to vary while maintaining the discount factor fixed at 0.9, we see that the policies are still effective as shown in Figure 5. Thus, changing the noise level in this case does not affect the effectiveness of the policies.

**Figure 5:**
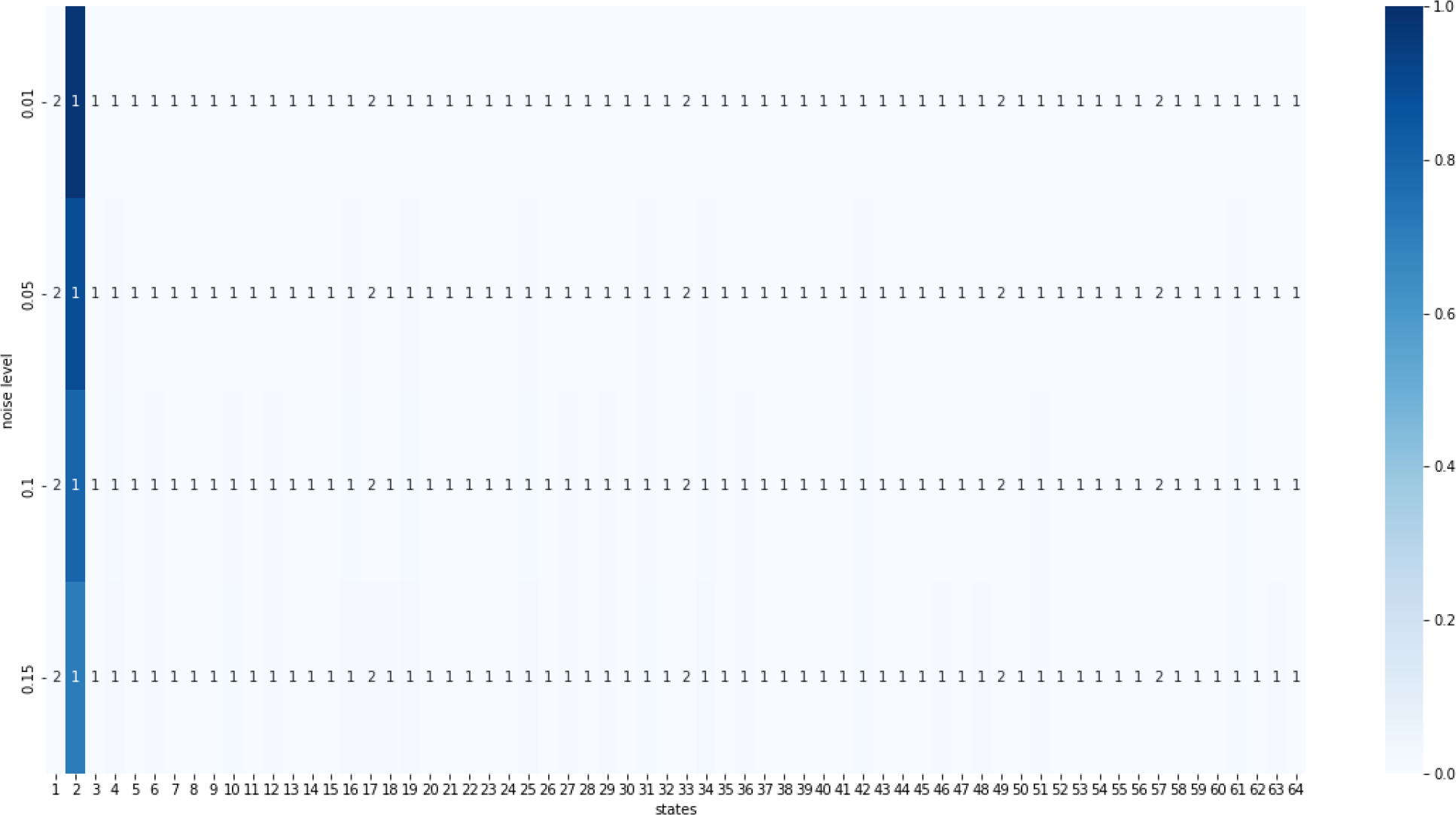
Heatmap of stationary distributions given all propensities are 0.9 using single-step optimal policies from the value iteration method. The states are on the *x*-axis and the *y*-axis gives the level of noise used to find the optimal policy. The discount factor was fixed to 0.9 for all policies, the stationary distribution consistently goes to state 2 (000001), so changing the noise in this case does not affect the effectiveness of the policies. The shading gradient on the right of the heatmap correspond to the proportion of time the process ends in one of the possible states.

**Figure 6:**
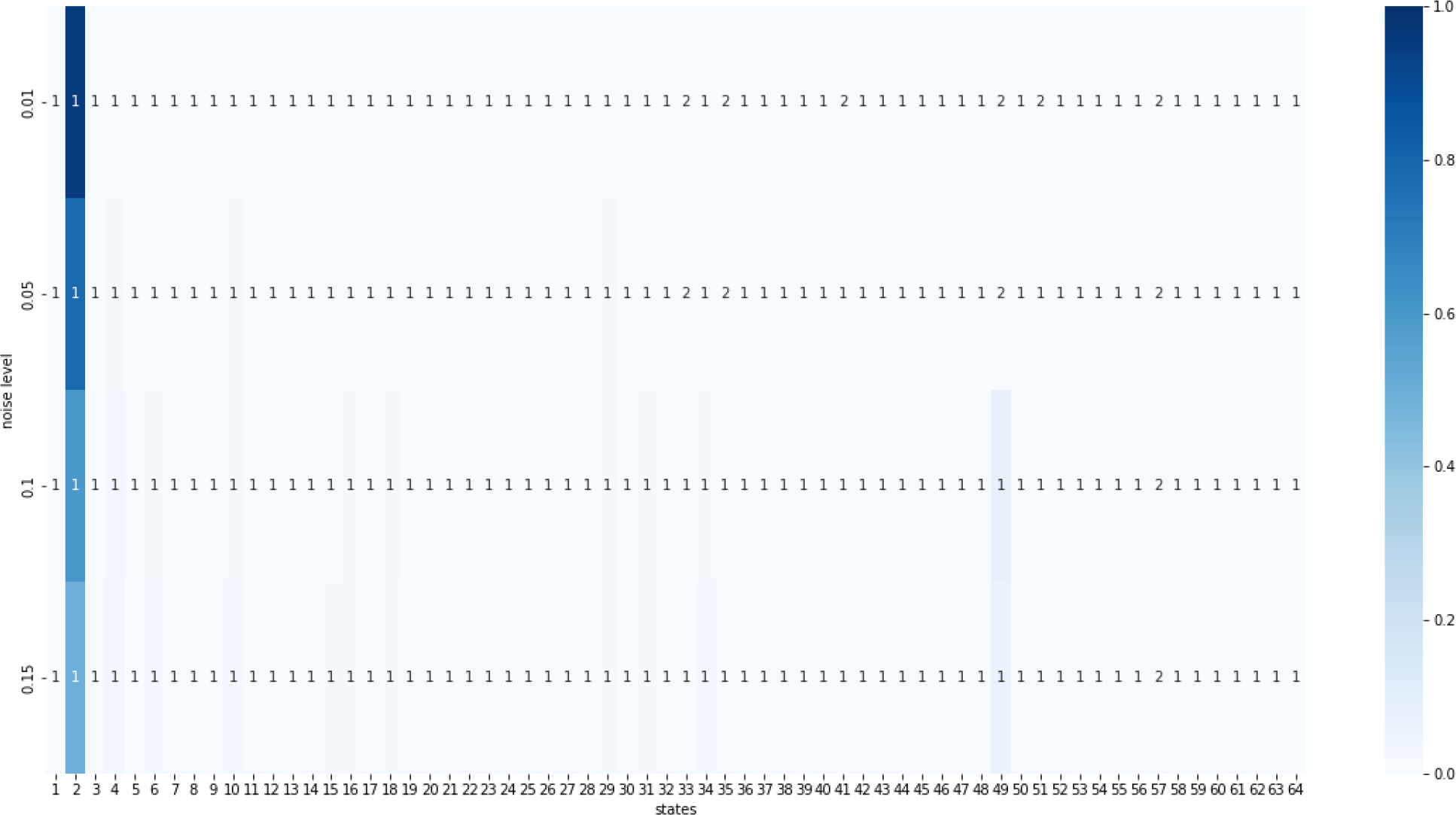
Heatmap of the stationary distribution given all propensities are 0.5 using single-step optimal policies from the value iteration method. The states are on the *x*-axis and the *y*-axis gives the level of noise used to find the optimal policy. The discount factor was fixed to 0.9 for all policies. For noise level of 0.05 or smaller, the stationary distribution consistently goes to state 2 (000001), so small levels of noise the policies becomes effective.

**Figure 7:**
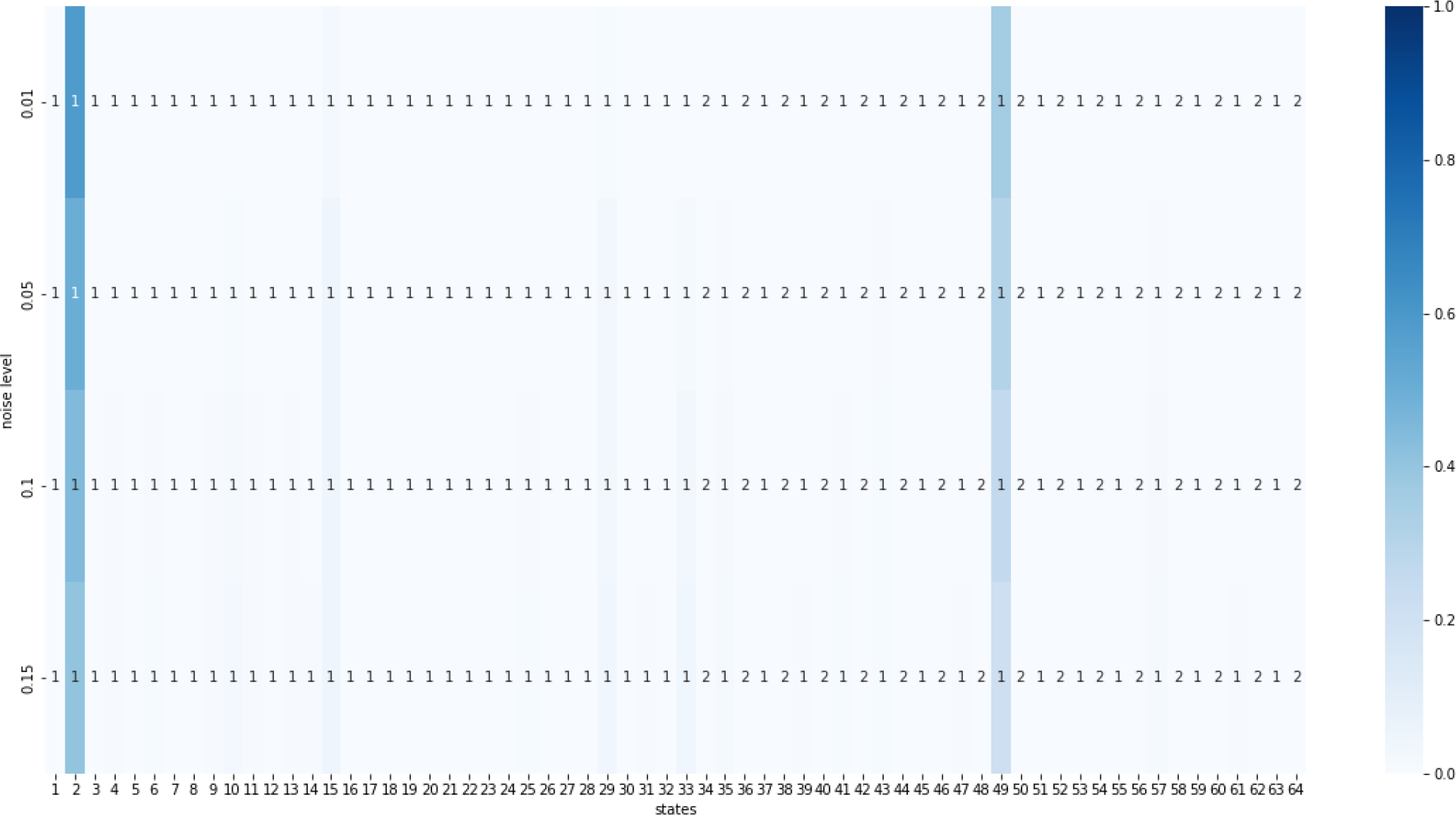
Heatmap of the stationary distribution given propensities from Table 1 using single-step optimal policies from the value iteration method. The states are on the *x*-axis and the *y*-axis gives the level of noise used to find the optimal policy. The policies for these choice of parameters are always ineffective, so changing the noise in this case does not affect the ineffectiveness of the policies.

In the second case, when all propensities are set to 0.5, allowing the noise level to vary while maintaining the discount factor fixed at 0.9, we see that some policies become effective while other are still ineffective as shown in Figure 6. Thus, changing the noise level in this case does affect the effectiveness of the policies. For noise level of 0.05 or smaller, the stationary distribution consistently goes to state 2 (000001), so for small levels of noise the policies becomes effective.

Finally, when we used propensities from Table 1, allowing the noise level to vary while maintaining the discount factor fixed at 0.9, we see that the policies for these choice of parameters are always ineffective. Thus, changing the noise level in this case does not affect the ineffectiveness of the policies.

We also computed optimal policies that have a duration period of *L* steps that are followed by a recovery period (without intervention) of *L − W* steps as described in [15, 16]. In Figures 15-17 we show results for optimal policies with *L* = *W* for *L* = 2, 4, 6, 8. We see that for the case of all propensities equal to 0.5, the policies become effective for a discount factor of 0.9 or higher. However, for the propensities in Table 1, the policies are still ineffective. For policies with four steps or higher, even the propensities in Table 1 will produce effective policies.

### 3.2 The Repressilator

The Repressilator is a gene regulatory network consisting of 3 genes, each expressing a protein that represses the next gene in the loop. Its wiring diagram is a cycle of three repressors as shown in Figure 8a. This model has been studied using differential equations [25]. The Repressilator has been used for the design and construction of synthetic networks to understand different cell functions [26]. Here we create a Boolean model for the Represillator using three update functions given in Figure 8b. Then we examine the effect the propensities have on its long term dynamics using the SDDS framework. The synchronous dynamics (i.e., without stochasticity) of this model is given by two cycles: a 2-cycle and a 6-cycle as shown in Figure 9a. We note that the 6-cycle represents a biological attractor that describes the stable oscillations of the three repressors are was shown to exist with other model types [26, 25]. The 2-cycle is an artificial attractor which a product of the synchronous update. Once we add stochasticity to the model using the SDDS framework, the states in the 2-cycle became transient states (that is, it was possible to escape from the 2-cycle). However, the 6-cycle was still an attractor as shown in Figure 9b. We highlight the fact that the biological attractor is robust to the update mode. Thus, we consider the six-cycle to be the absorbing state for calculating time to absorption. We note that for this example we do not consider control actions and therefore this example is not a Markov decision process. However, we will study the effect of changes in the propensity parameters on the time to absorption and the mixing time of this model.

**Figure 8:**
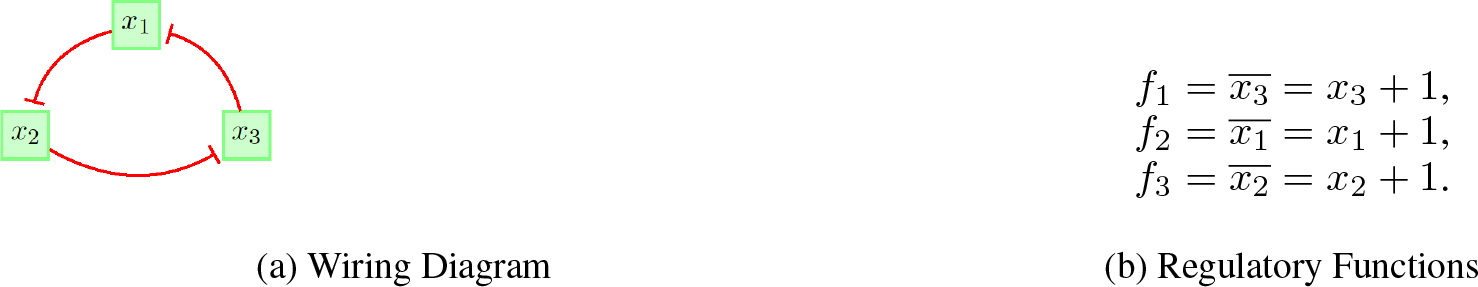
Boolean model of the Represillator.

**Figure 9:**
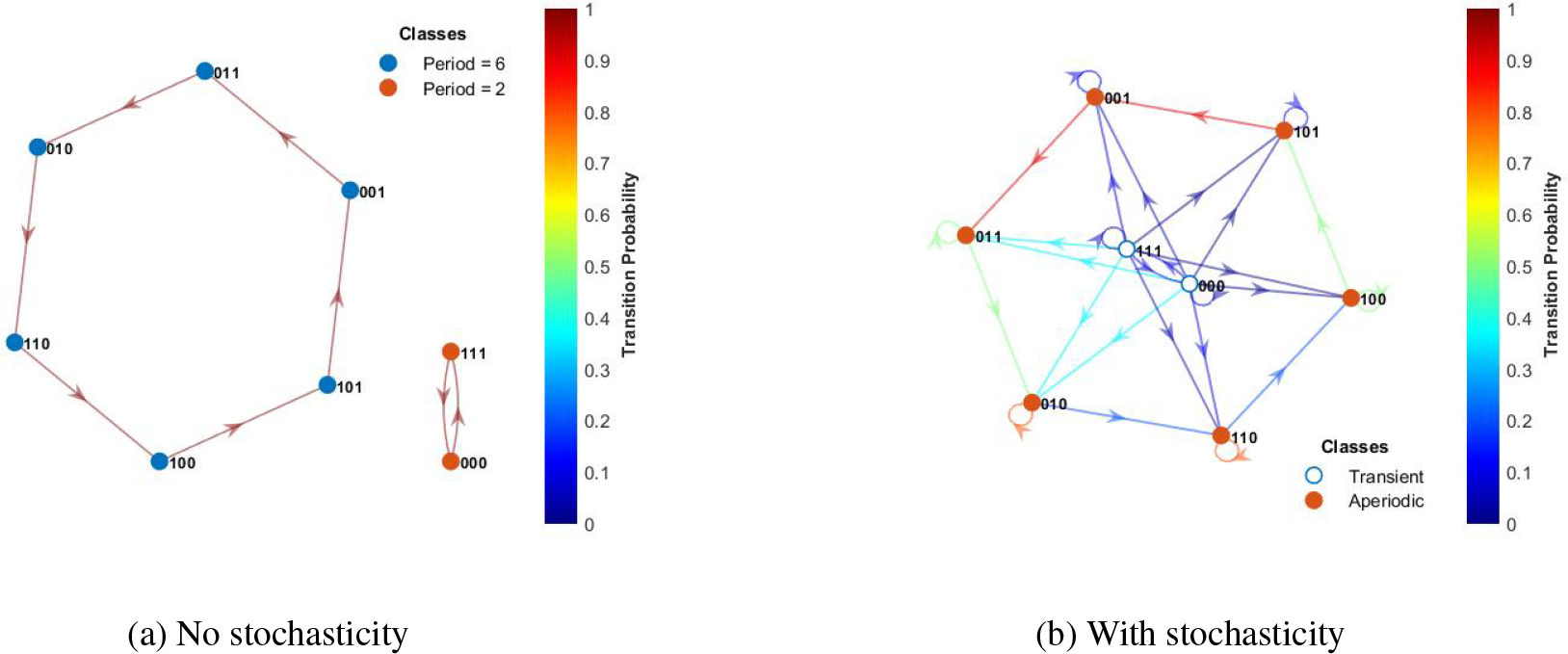
The synchronous dynamics are given by a 6-cycle and a 2-cycle in 9a without noise. With stochasticity using the SDDS framework, one may leave the two-cycle for the six-cycle, but not the other way around.

In Figure 9b we can see the effect the propensities have on the transition from one state to another. In this particular example the propensities are 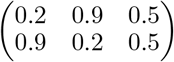, where the top row is the activation propensity for each function and the bottom row is the degradation propensity for each function. We note that in this selection of propensities we are specifying a slow activation rate for the variable *x*_1_ and a fast degradation rate for variable *x*_1_. Inversely, we are specifying a fast activation rate for the variable *x*_2_ and a slow degradation rate for variable *x*_2_. The mixing time is 4.5547 steps with the repressilator spending 6.85% of the time in each state 001 and 101, 12.33% of the time in each state 011 and 100, and 30.82% of the time in each state 010 and 110. Negligible time is spent in states 000 and 111. We want to investigate the range of mixing times for the Repressilator.

First, we examine the mixing time when both propensities for all variables are the same. In Figure 10, the top line in blue represents the mixing times when all propensities are the same. We find the lowest mixing time when all of the propensities are 0.5. The red line represents when we only change the propensities inversely for 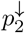 and 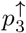 while keeping all other propensities 0.5. In this case, we find that the mixing time is lowest at the ends of the graph. That is when 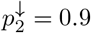 and 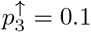 and vice versa, we achieve the lowest mixing time.

**Figure 10:**
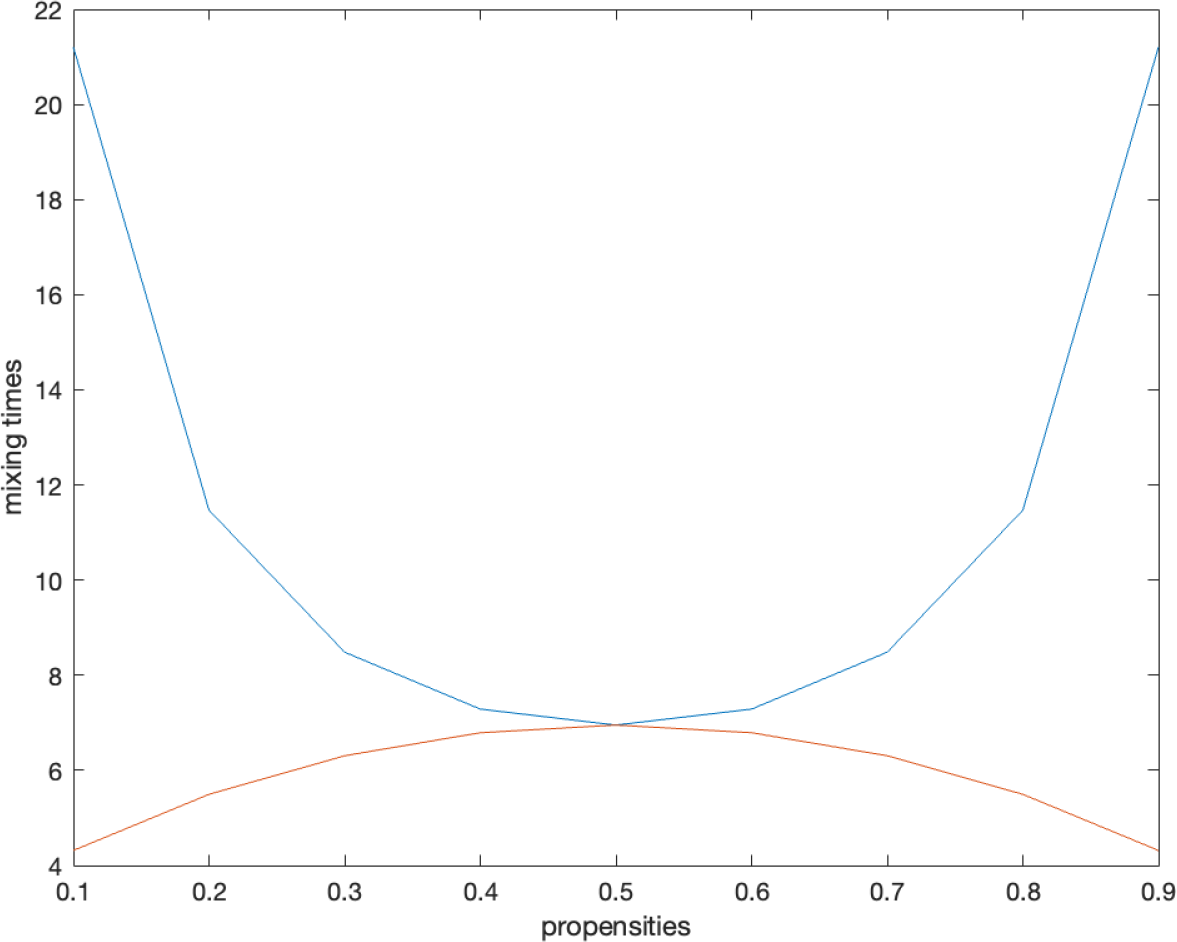
A comparison of the mixing times. The *x*-axis represents the value of the propensities. The *y*-axis represents the mixing time. The blue line represents the mixing times when all propensities are the same. We find the lowest mixing time when all of the propensities are 0.5. The red line represents when we only change the propensities inversely for 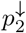 and 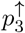 while keeping all other propensities 0.5. In this case, we find that the mixing time is lowest at the ends of the graph. That is when 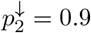 and 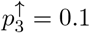 and vice versa, we achieve the lowest mixing time.

It is intuitive that when all of the propensities are low, the states in the Repressilator are less likely to change their current states. In particular, since the two-cycle is less likely to change states, the time to absorption becomes larger which causes the mixing time to be higher as well (see Figure 11a). A similar effect is seen with high propensities where the 000 and 111 alternate before being absorbed as in Figure 11b. The smallest mixing time appears to occur when the propensities are at 0.5 when the system has the highest change to leave the two-cycle and be absorbed (Figure 12a). However this mixing time is 6.9521, which is higher than the mixing time for the starting example. The minimum mixing time seems to occur when the propensities are not equal.

**Figure 11:**
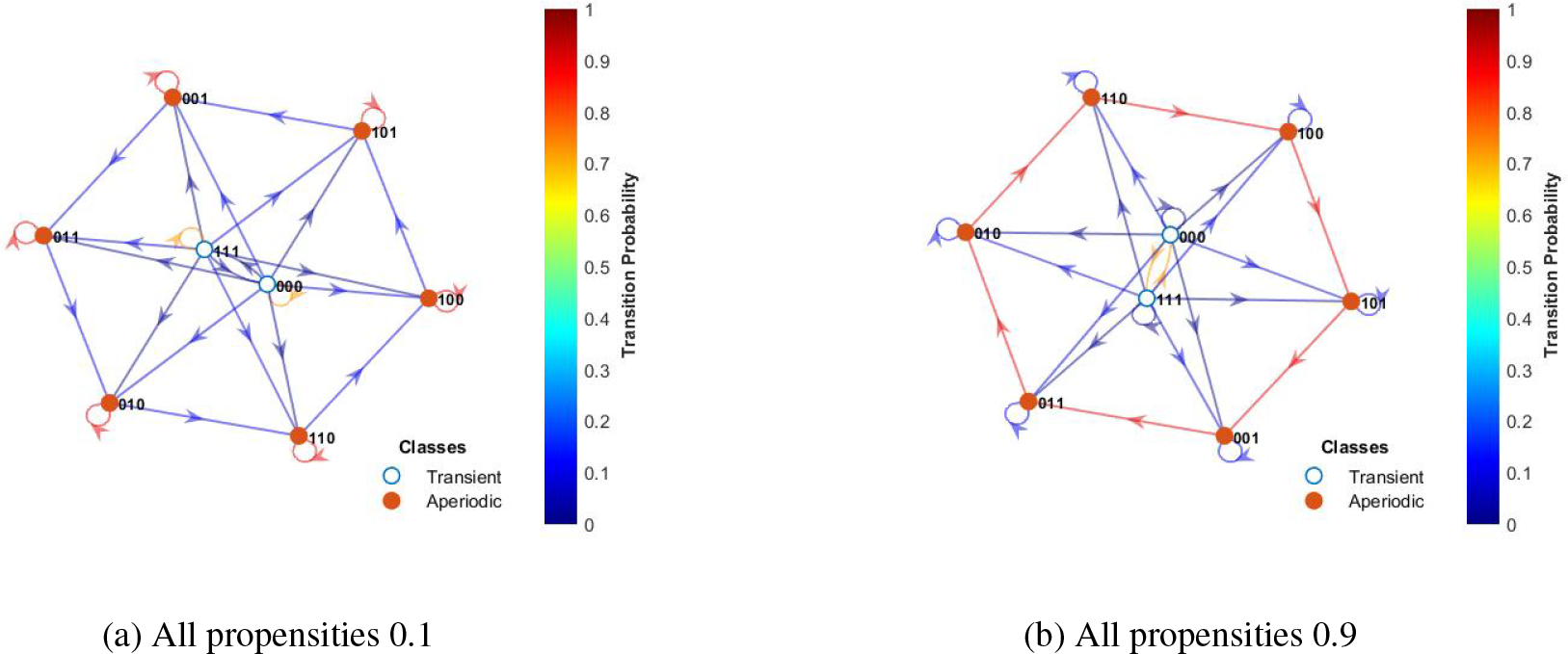
Same propensities comparison: When the propensities are all 0.1, the network has a tendency to stay at the state it started on and when the propensities are all 0.9, it has a tendency to go to the next state in the cycle more quickly. In both cases, the probability to leave the two-cycle is low.

**Figure 12:**
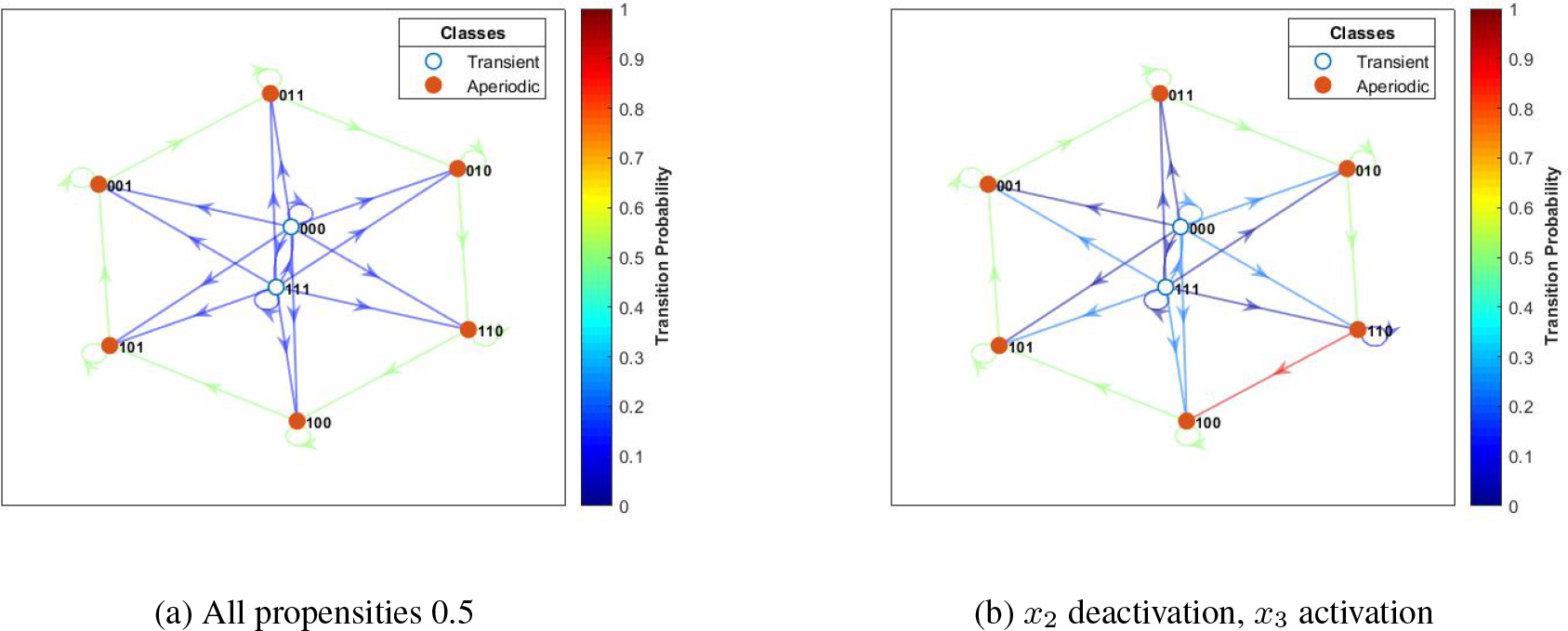
When all of the propensities are 0.5, the system has the fastest mixing time of all same propensities. When 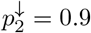 and 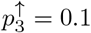, we see that state 001 has a much higher probability to go to state 100 in comparison to all other probabilities in the network. This is when the mixing time is fastest overall.

To further analyze the effect of changes in propensities, we start by setting all propensities to 0.5 and allowing one propensity to differ. The lowest mixing time for propensities tested from 0.1 to 0.9 is 4.8887 at 0.1 and remains the same for all variables. Allowing two functions to have differing inverse propensities yields more interesting results. For example, if the propensities are 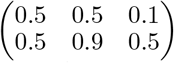, we get a mixing time of 4.3157. The mixing times for propensities for this inverse relationship between the activation propensity for the third function and the deactivation propensity for the second function can be seen in Figure 12b starting at 0.1 and 0.9 respectively. Setting all other propensities to 0.9 allows an even shorter mixing time of 2.8627 at endpoints 0.1 and 0.9, but setting all other propensities to 0.1 forces the mixing time to be even longer with 11.9877 being the smallest mixing time at 0.5. Finally, it is worth noting that the time to absorption remains low even when mixing times are large. For instance, when the mixing time is 11.8977 for propensity matrix 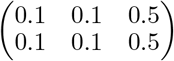, the time to absorption is 1.6949. In summary, we have observed that the mixing times of this model are sensitive to changes in the propensity parameters while the time to absorption remained stably low. This means that it is always possible to escape from the artificial attractor and then to transition into the biologically-relevant attractor relatively quickly, and after a number of steps given by the mixing time the stable oscillations will be achieved.

We show the transition matrix (obtained using Eq. 2) of the Boolean Repressilator model in Figure 13 and their eigenvalues in Equation 13. When all the propensities are the same (i.e., *p* = 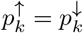 for all *k*), we see that the absolute value of all eigenvalues are minimized when 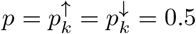 as shown in Figure 14.

**Figure 13:**
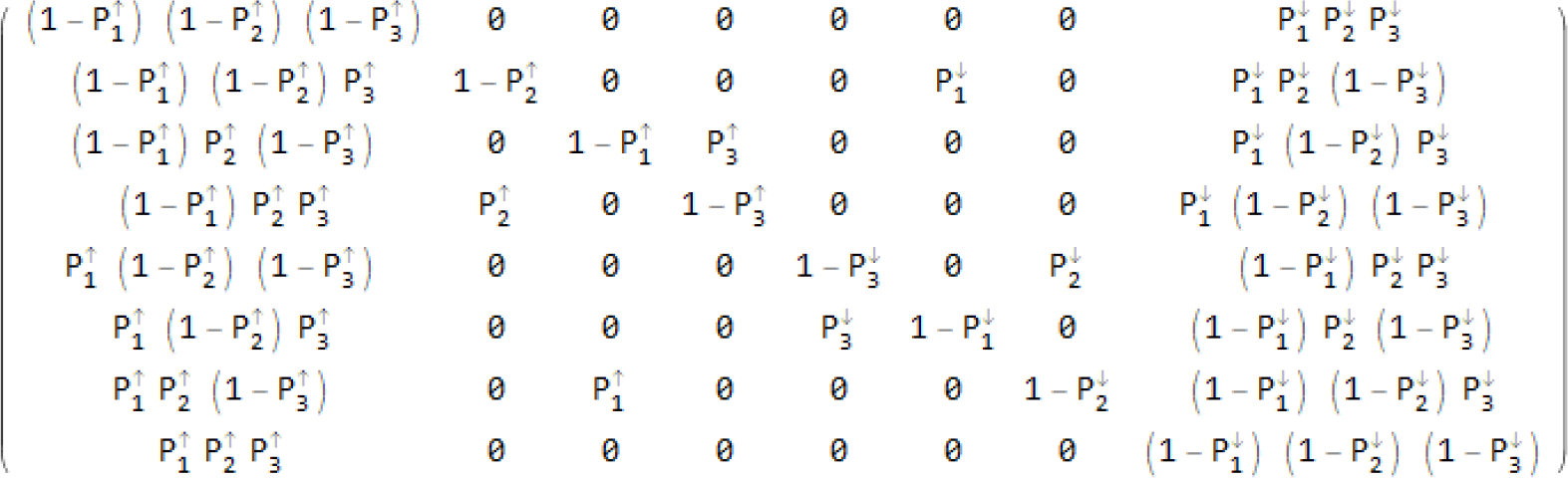
Transition matrix of the Repressilator model.

**Figure 14:**
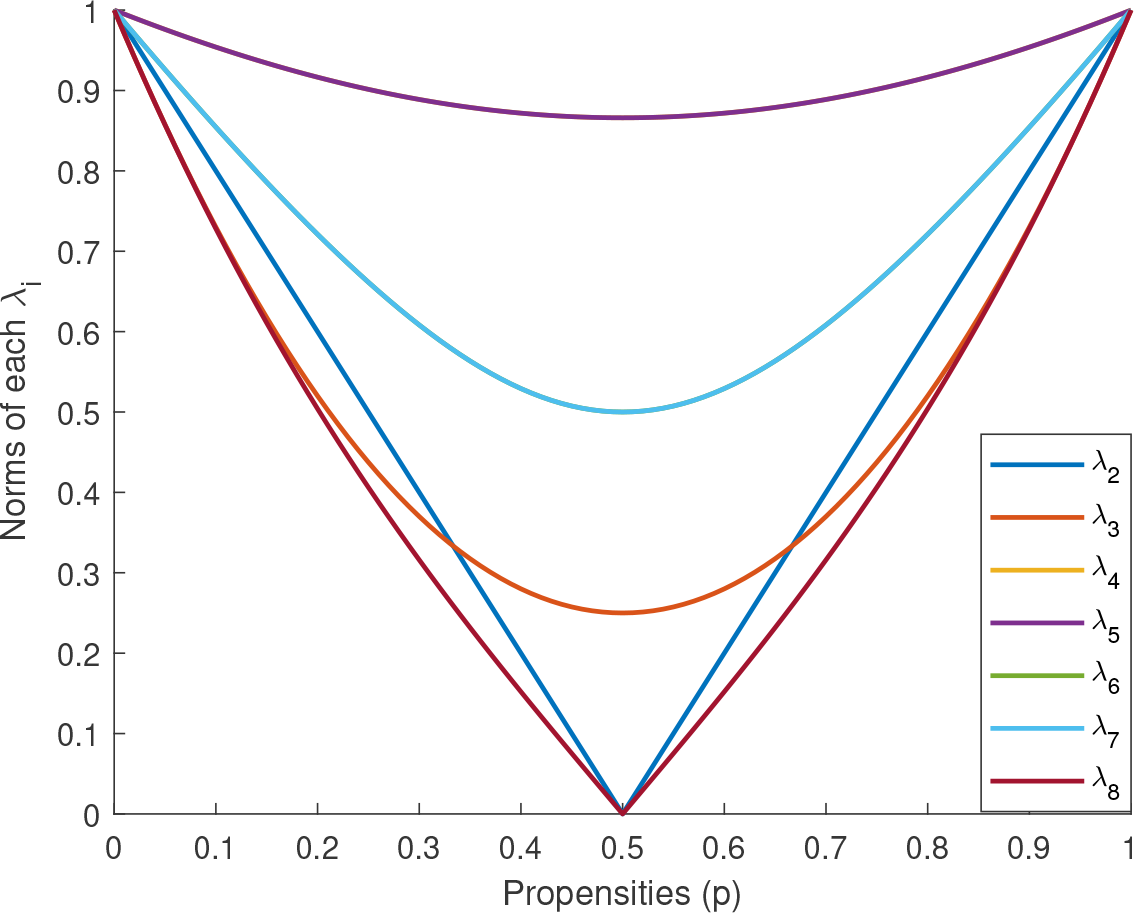
Here we show the plots of eigenvalues from the transition matrix in Figure 13, where all propensities are assumed to be the same. Eigenvalues (listed in order in Equation 13) are all minimized when the propensity is *p* = 0.5, and graphical results can easily be confirmed analytically or numerically with MATLABs *fminsearch* function.

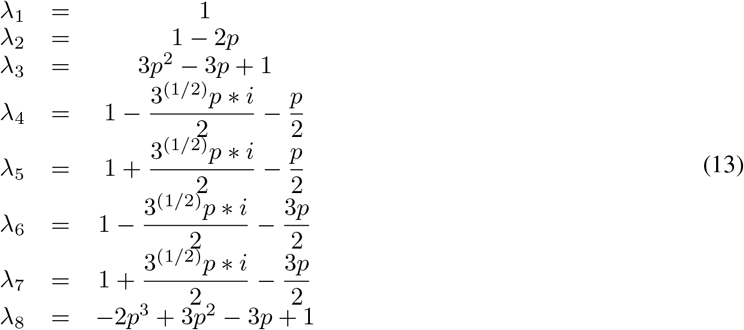

## 4 Conclusion

In this paper, we studied the effect of changes in model parameters on optimal control policies, time to absorption, and mixing times by focusing on a class of Markov decision processes that is obtained by considering stochastic Boolean networks equipped with control actions. The stochastic framework that we used is the one called Stochastic Discrete Dynamical Systems (SDDS). The SDDS framework introduces stochasticity by assigning two propensity parameters to each function in a Boolean network. Using the SDDS framework and their control actions, we have performed an analysis of the effect of changes in propensity parameters on optimal control policies, time to absorption, and mixing times on specific Boolean models.

In the first part of the results, we studied how the choice of propensity parameters affects the effectiveness of optimal control policies where, depending on the range of values of certain parameters (e.g., the discount factor, level of noise, and the period of control policy duration), some policies became ineffective. We have also shown that by tuning the noise level and duration of the control policies one can still guarantee the effectiveness of some of the policies that we have considered.

In the second part of the results, we have studied the effect of changes in the propensity parameters on the time to absorption and the mixing time for a Boolean model of the Repressilator. We have shown that the mixing time of Repressilator model was very sensitive to changes in the propensity parameters while the time to absorption remained stably low. This means that in this model, regardless of the propensities, it will always be possible to reach the biologically-meaningful attractor relatively quickly, and after a number of steps specified by the mixing time the stable oscillations will be achieved.

To conclude, the results of this paper will help us understand the sensitivities of model parameters in similar models and their results and might help us inform the selection of different parameters in other models.

## Appendix

In this section we provide additional heatmaps for multi-period of optimal control policies for the T-LGL model discussed in Section 3.1.

**Figure 15:**
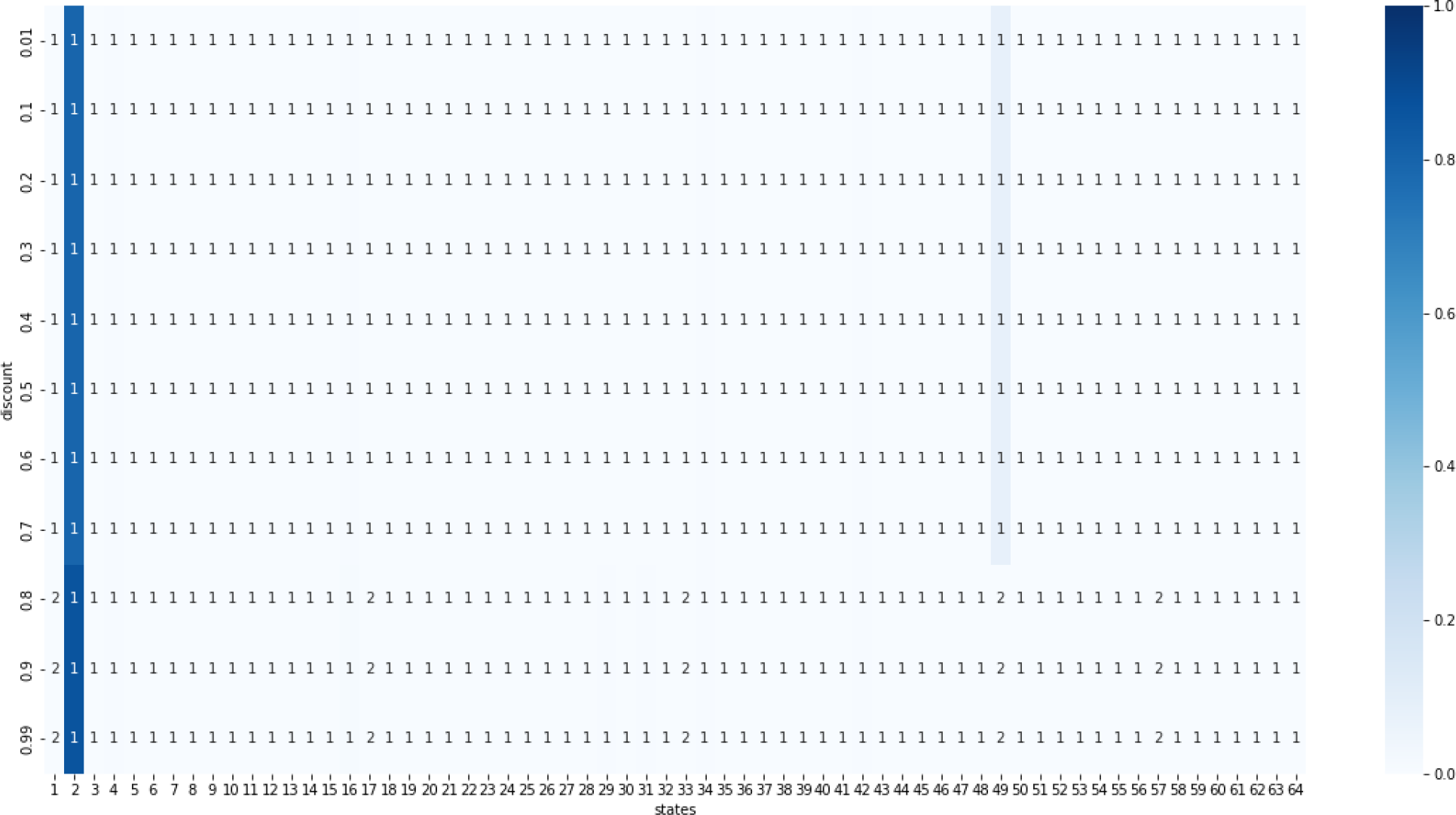
Heatmap of stationary distributions given all propensities are 0.9 using two-step optimal policies from the value iteration method. The states are on the *x*-axis and the *y*-axis gives the discount used to find the policy. At a discount of 0.8 or higher, the stationary distribution consistently goes to state 2 (000001), so this is when the policy becomes effective.

**Figure 16:**
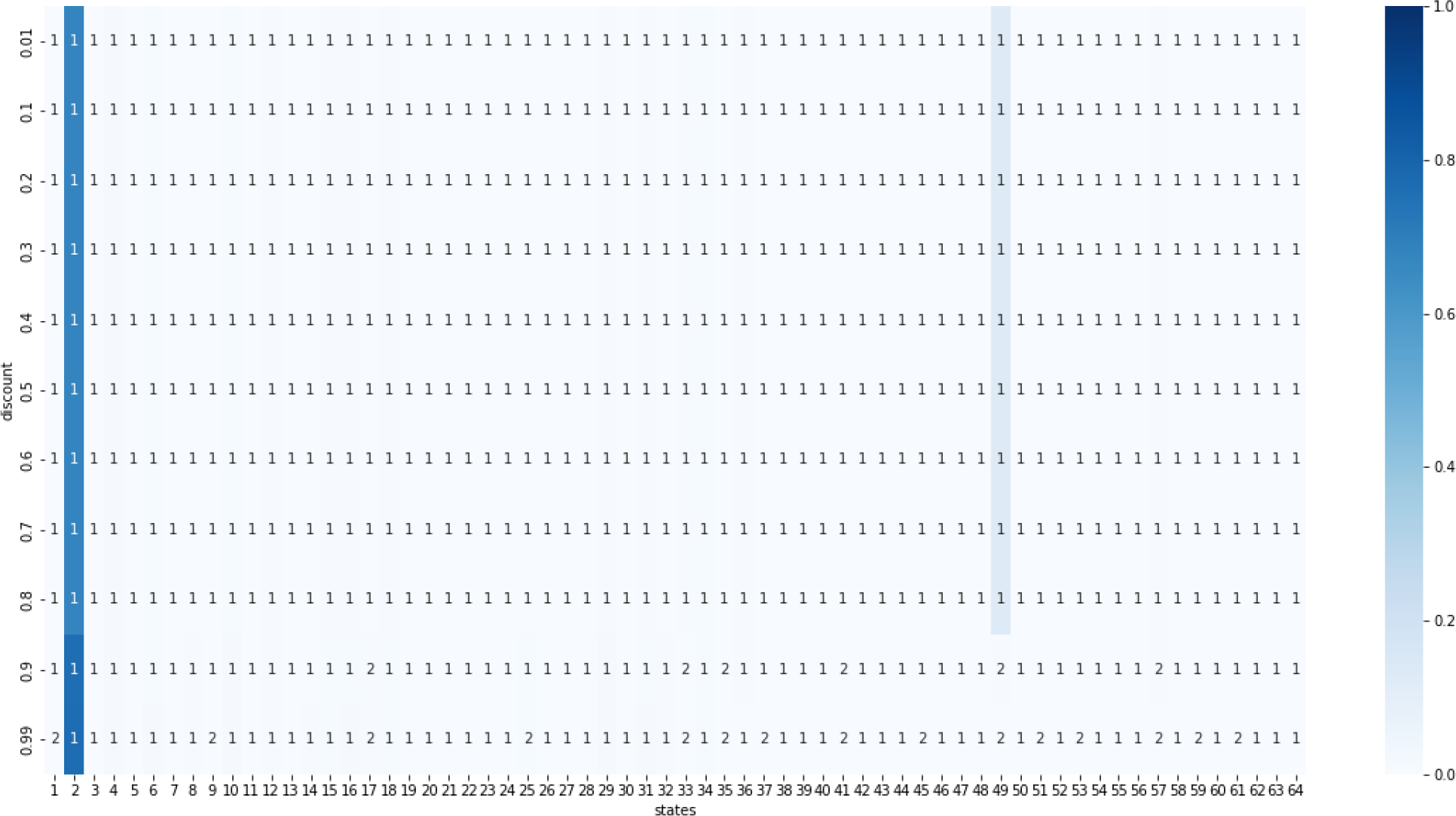
Heatmap of the stationary distribution given all propensities are 0.5 using two-step optimal policies from the value iteration method. The states are on the *x*-axis and the *y*-axis gives the discount used to find the policy. At a discount of 0.9 or higher, the stationary distribution consistently goes to state 2 (000001), so this is when the policy becomes effective.

**Figure 17:**
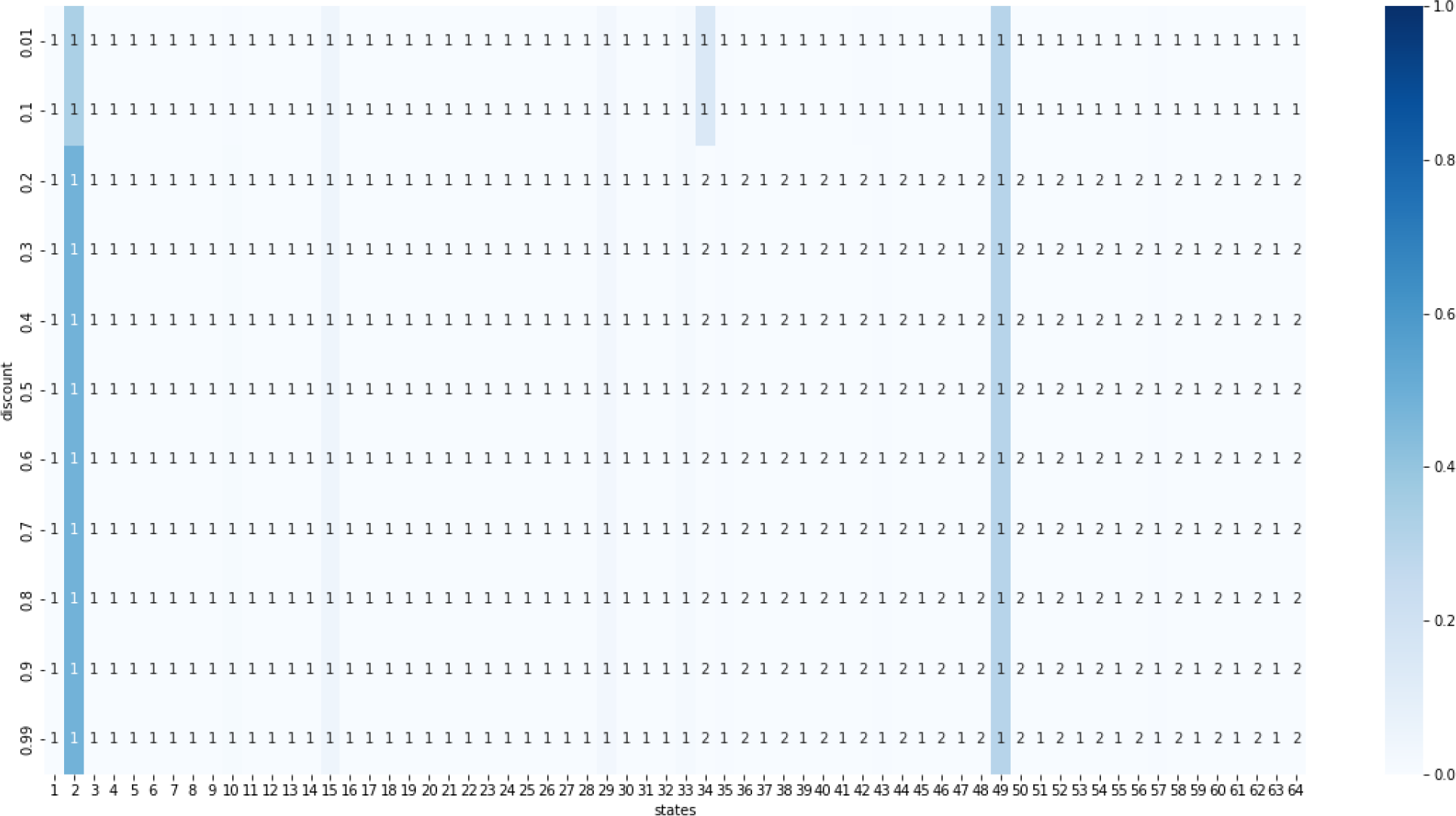
Heatmap of the stationary distribution given propensities from Table 1 using two-step optimal policies from the value iteration method. The states are on the *x*-axis and the *y*-axis gives the discount used to find the policy. The policies for these choice of parameters are always ineffective.

**Figure 18:**
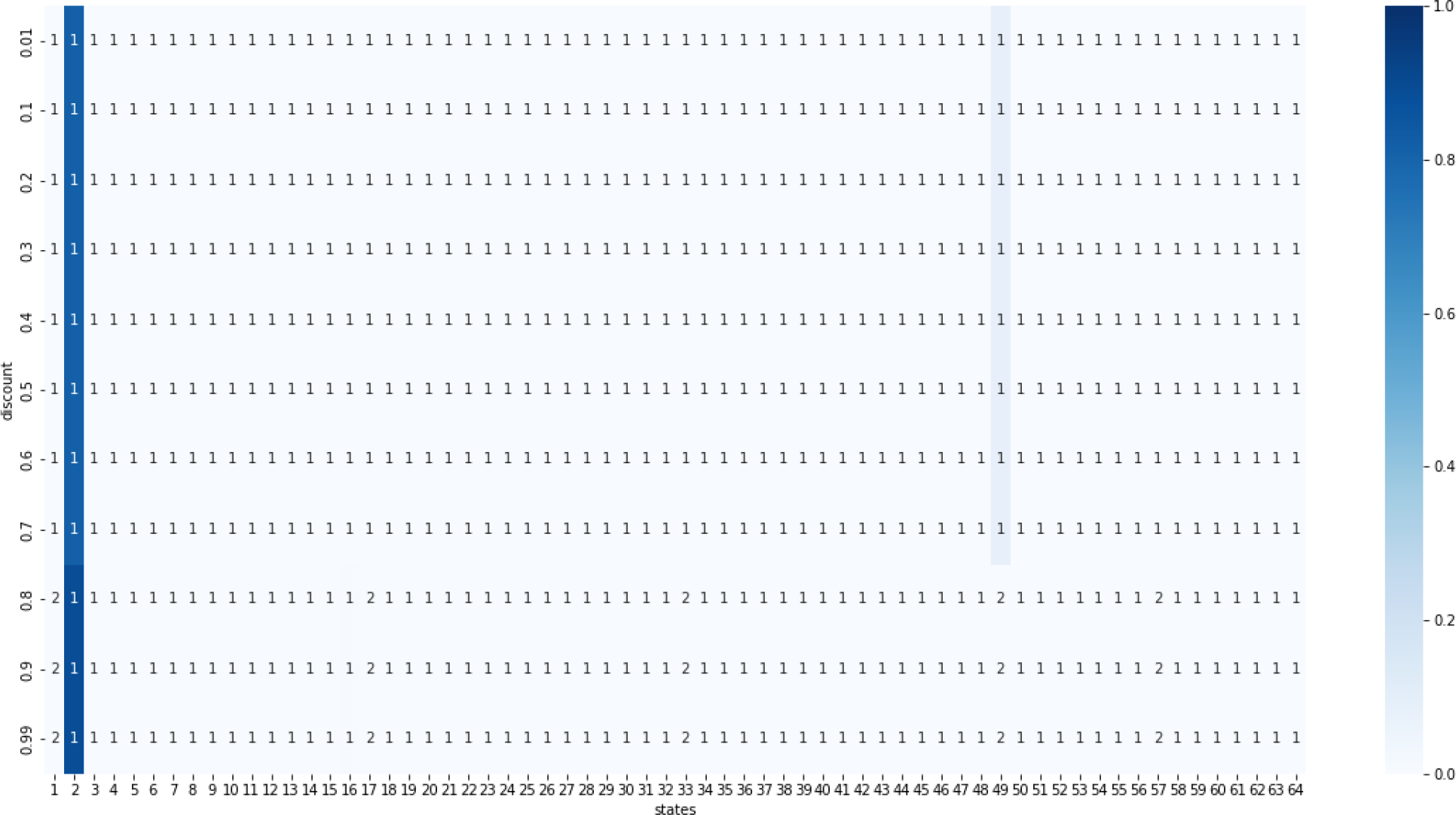
Heatmap of stationary distributions given all propensities are 0.9 using four-step optimal policies from the value iteration method. The states are on the *x*-axis and the *y*-axis gives the discount used to find the policy. At a discount of 0.8 or higher, the stationary distribution consistently goes to state 2 (000001), so this is when the policy becomes effective.

**Figure 19:**
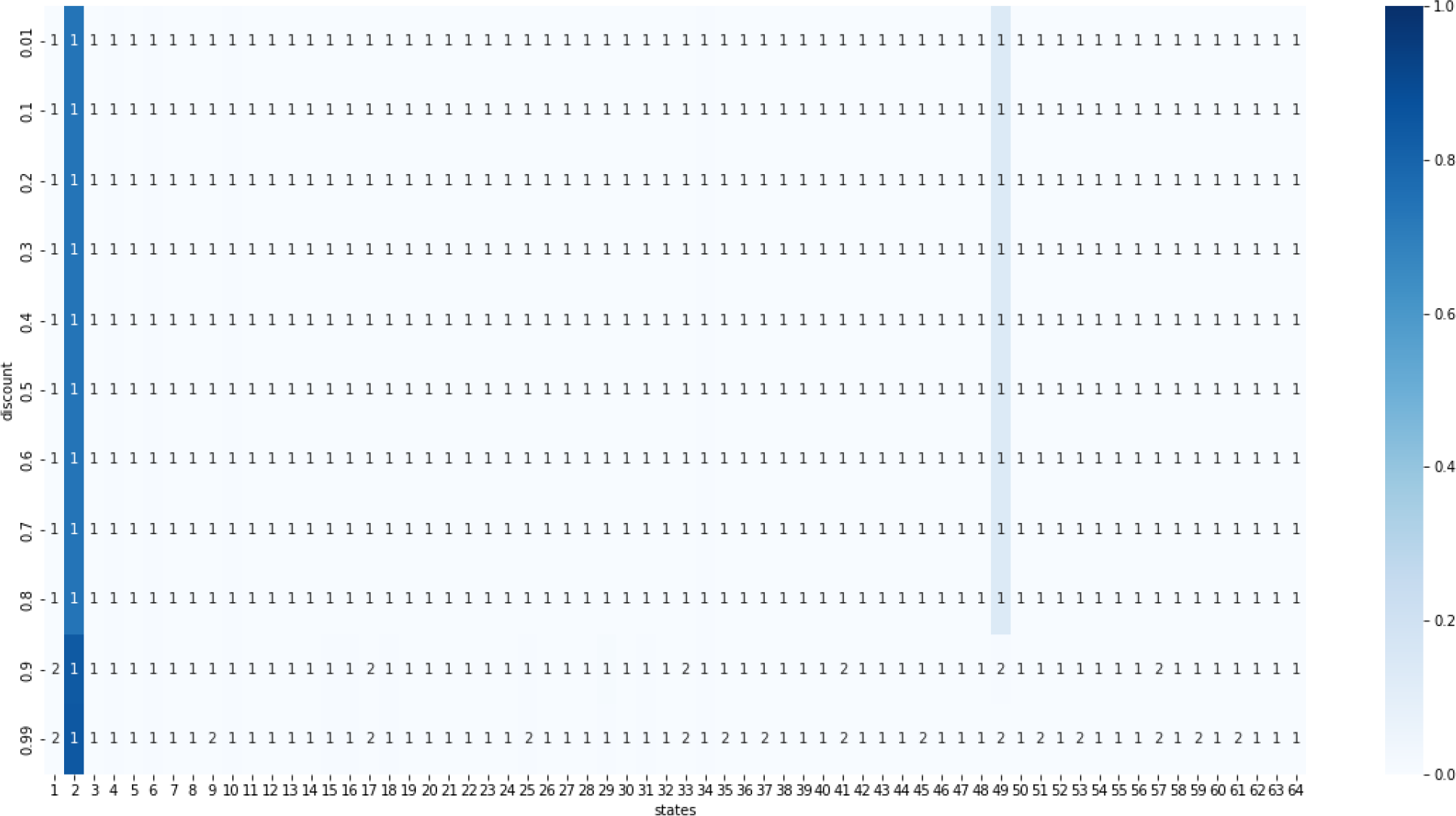
Heatmap of the stationary distribution given all propensities are 0.5 using four-step optimal policies from the value iteration method. The states are on the *x*-axis and the *y*-axis gives the discount used to find the policy. At a discount of 0.9 or higher, the stationary distribution consistently goes to state 2 (000001), so this is when the policy becomes effective.

**Figure 20:**
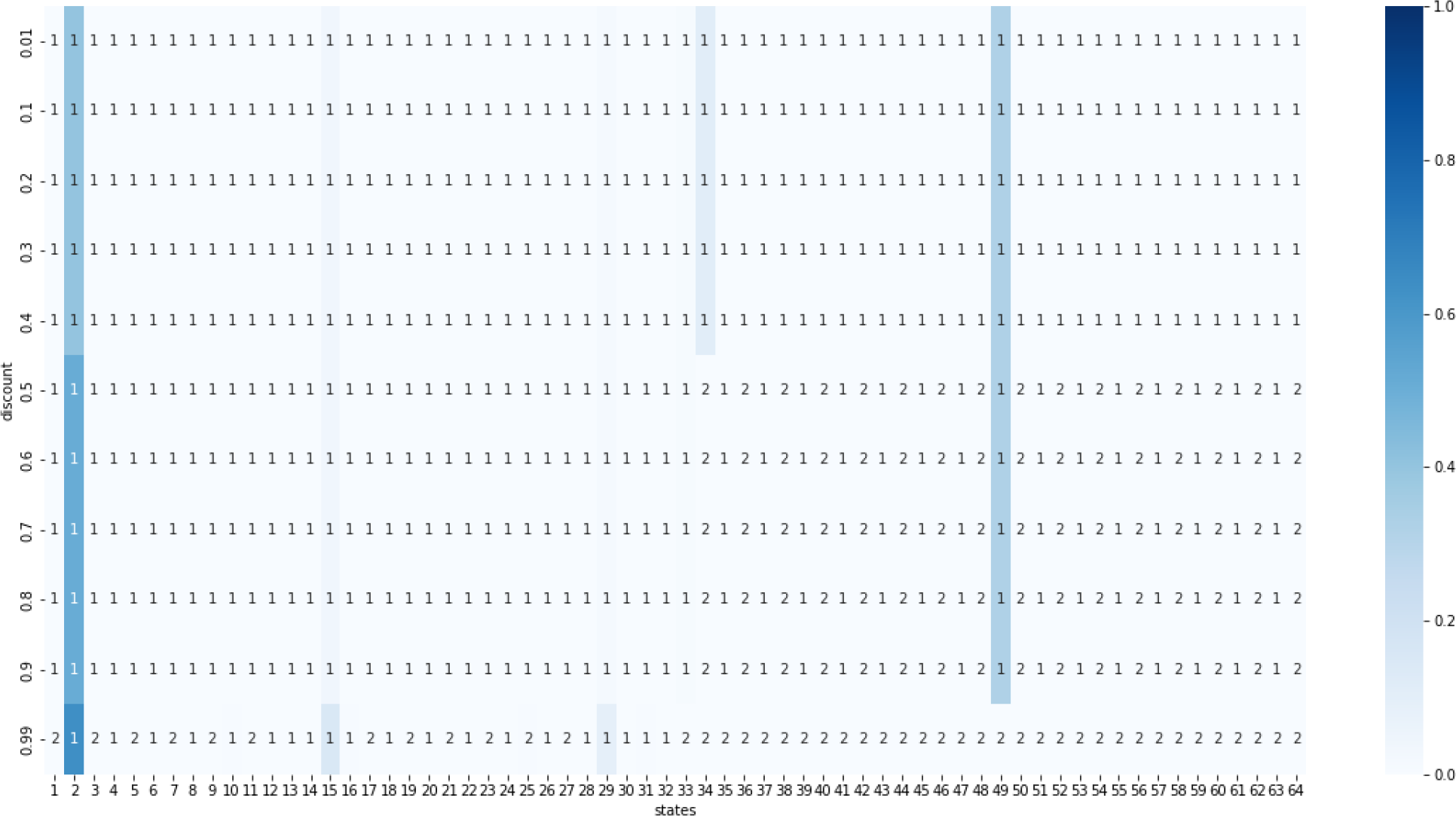
Heatmap of the stationary distribution given propensities from Table 1 using four-step optimal policies from the value iteration method. The states are on the *x*-axis and the *y*-axis gives the discount used to find the policy. The policies for these choice of parameters are always ineffective.

**Figure 21:**
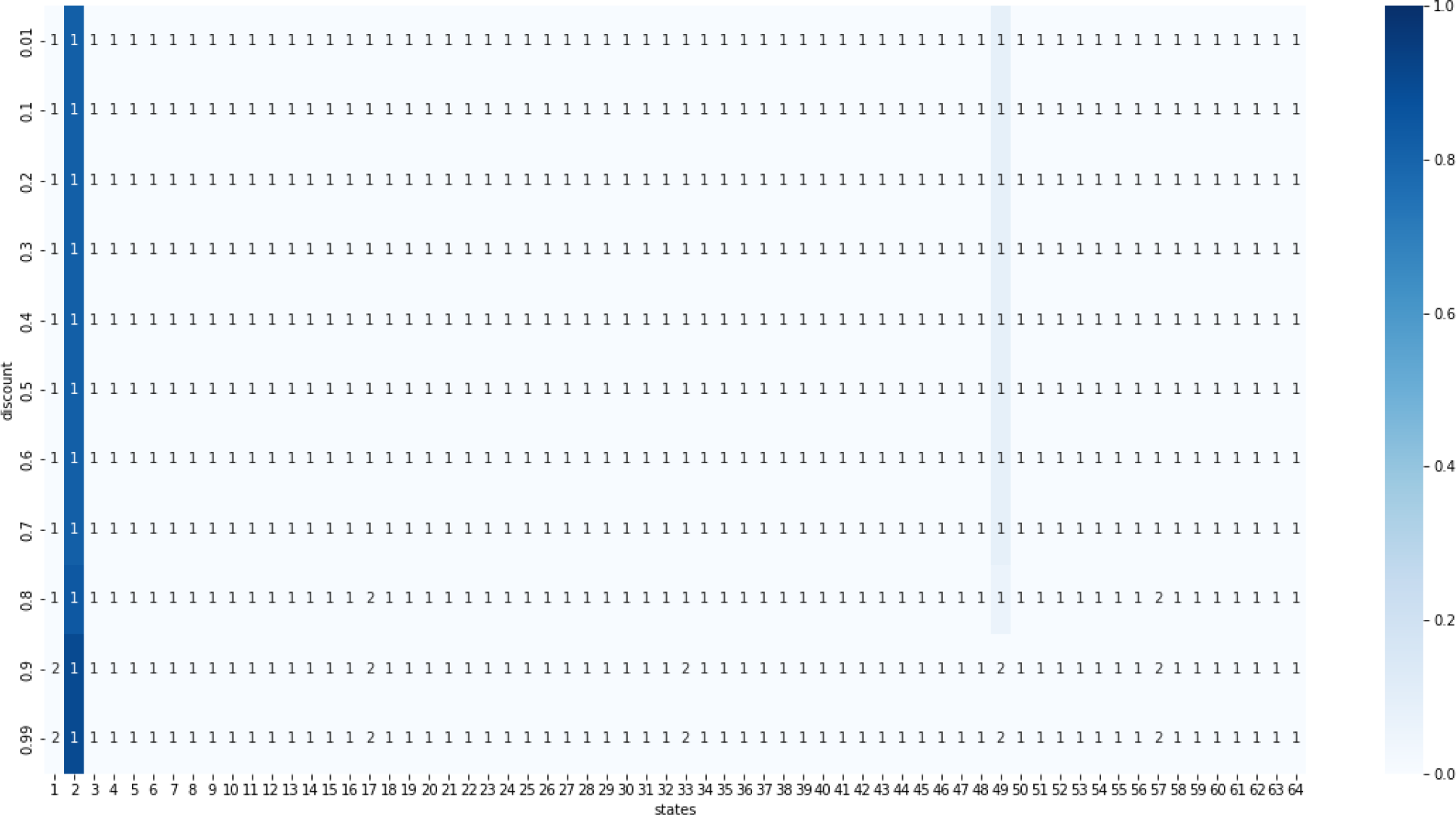
Heatmap of stationary distributions given all propensities are 0.9 using six-step optimal policies from the value iteration method. The states are on the *x*-axis and the *y*-axis gives the discount used to find the policy. At a discount of 0.8 or higher, the stationary distribution consistently goes to state 2 (000001), so this is when the policy becomes effective.

**Figure 22:**
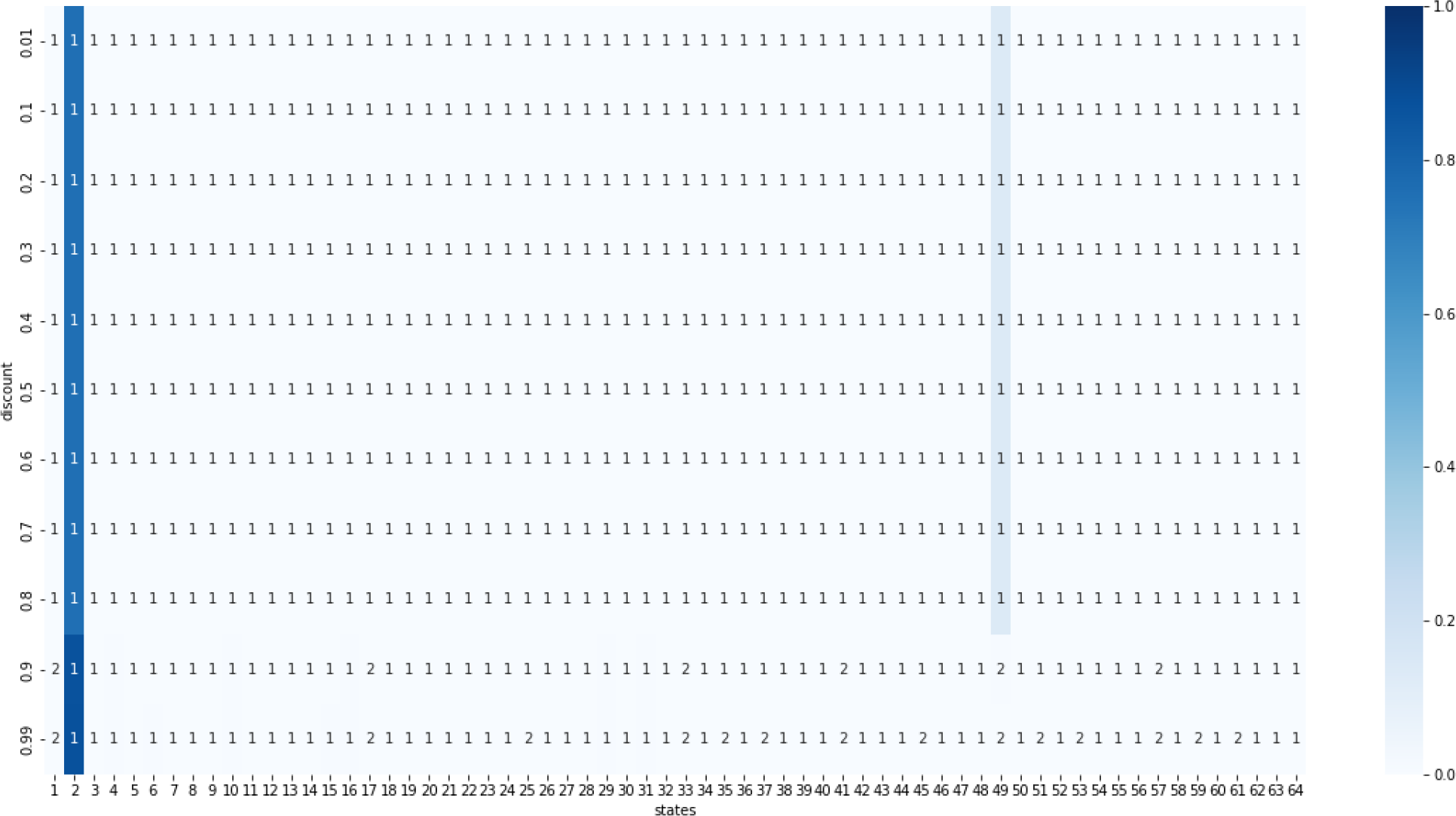
Heatmap of the stationary distribution given all propensities are 0.5 using six-step optimal policies from the value iteration method. The states are on the *x*-axis and the *y*-axis gives the discount used to find the policy. At a discount of 0.9 or higher, the stationary distribution consistently goes to state 2 (000001), so this is when the policy becomes effective.

**Figure 23:**
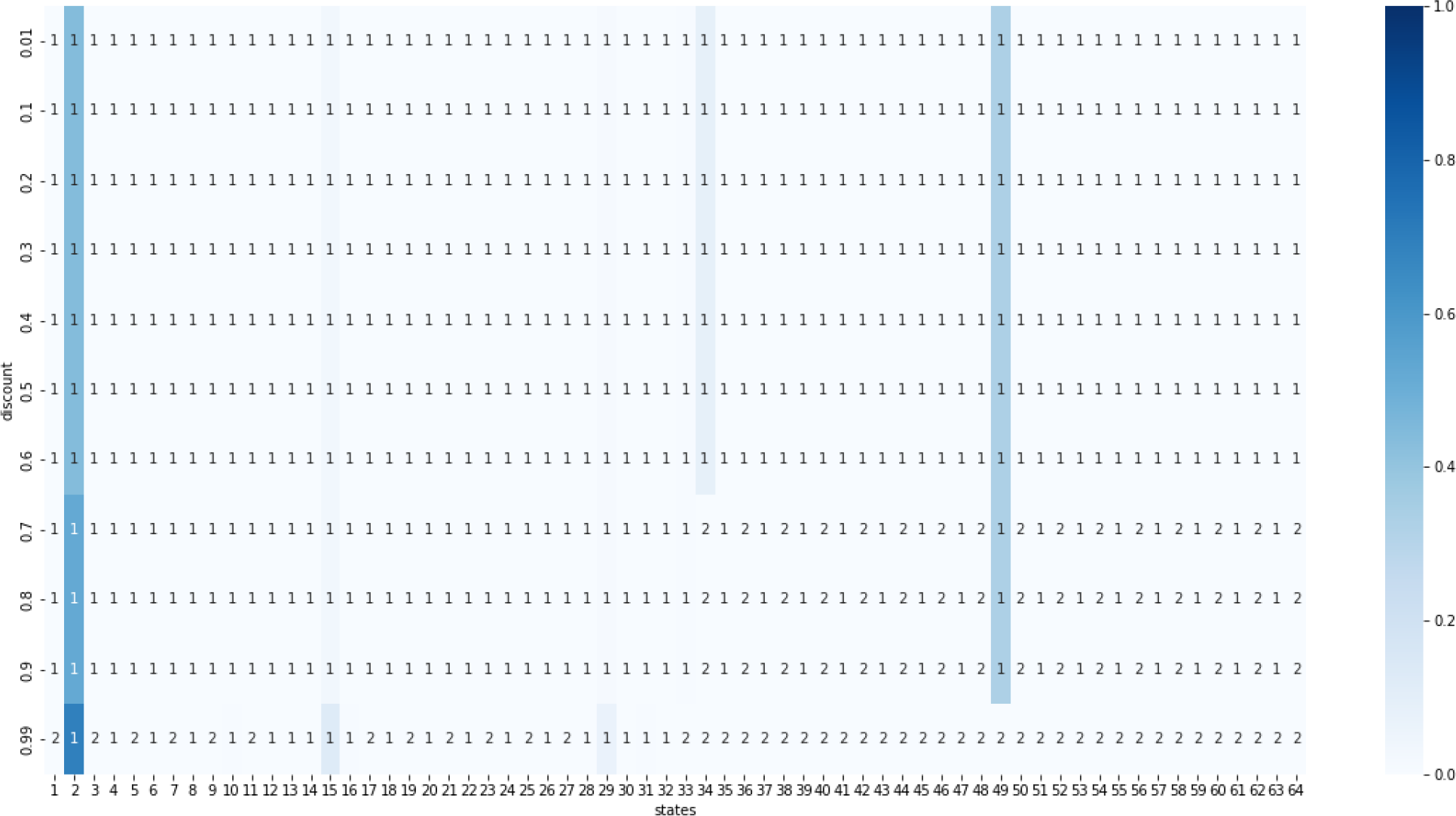
Heatmap of the stationary distribution given propensities from Table 1 using six-step optimal policies from the value iteration method. The states are on the *x*-axis and the *y*-axis gives the discount used to find the policy. The policies for these choice of parameters are always ineffective.

**Figure 24:**
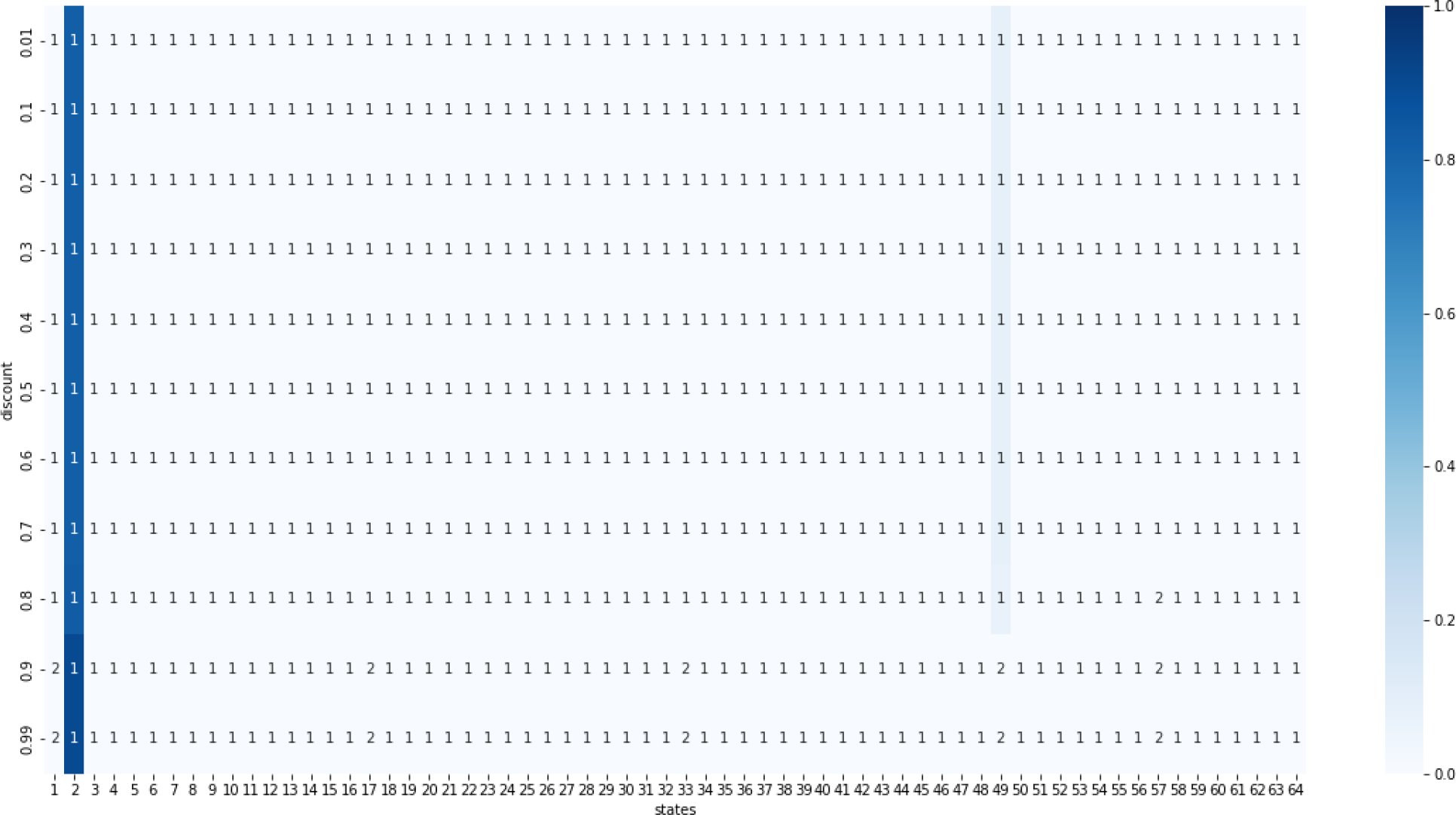
Heatmap of stationary distributions given all propensities are 0.9 using eight-step optimal policies from the value iteration method. The states are on the *x*-axis and the *y*-axis gives the discount used to find the policy. At a discount of 0.8 or higher, the stationary distribution consistently goes to state 2 (000001), so this is when the policy becomes effective.

**Figure 25:**
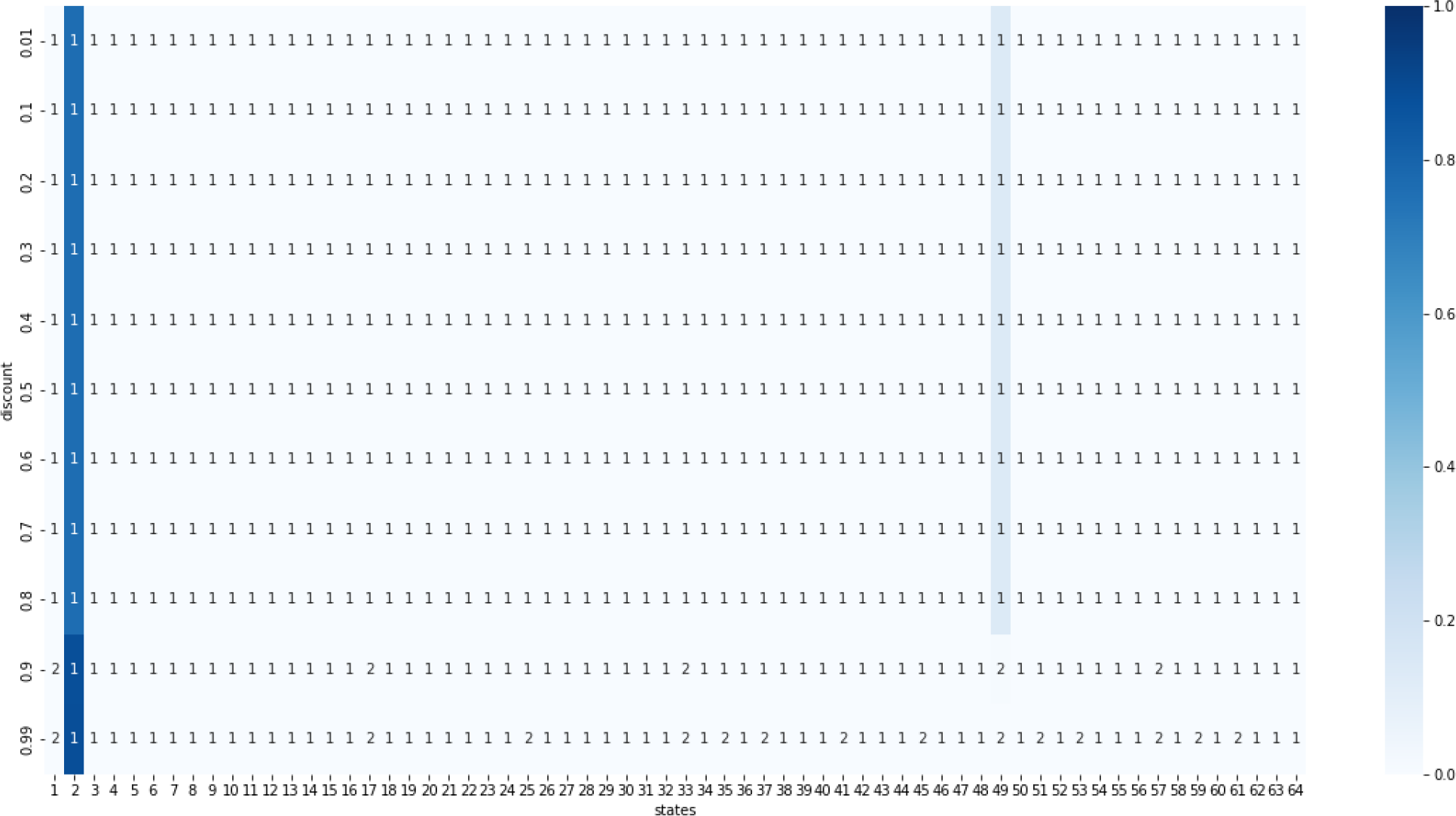
Heatmap of the stationary distribution given all propensities are 0.5 using eight-step optimal policies from the value iteration method. The states are on the *x*-axis and the *y*-axis gives the discount used to find the policy. At a discount of 0.9 or higher, the stationary distribution consistently goes to state 2 (000001), so this is when the policy becomes effective.

**Figure 26:**
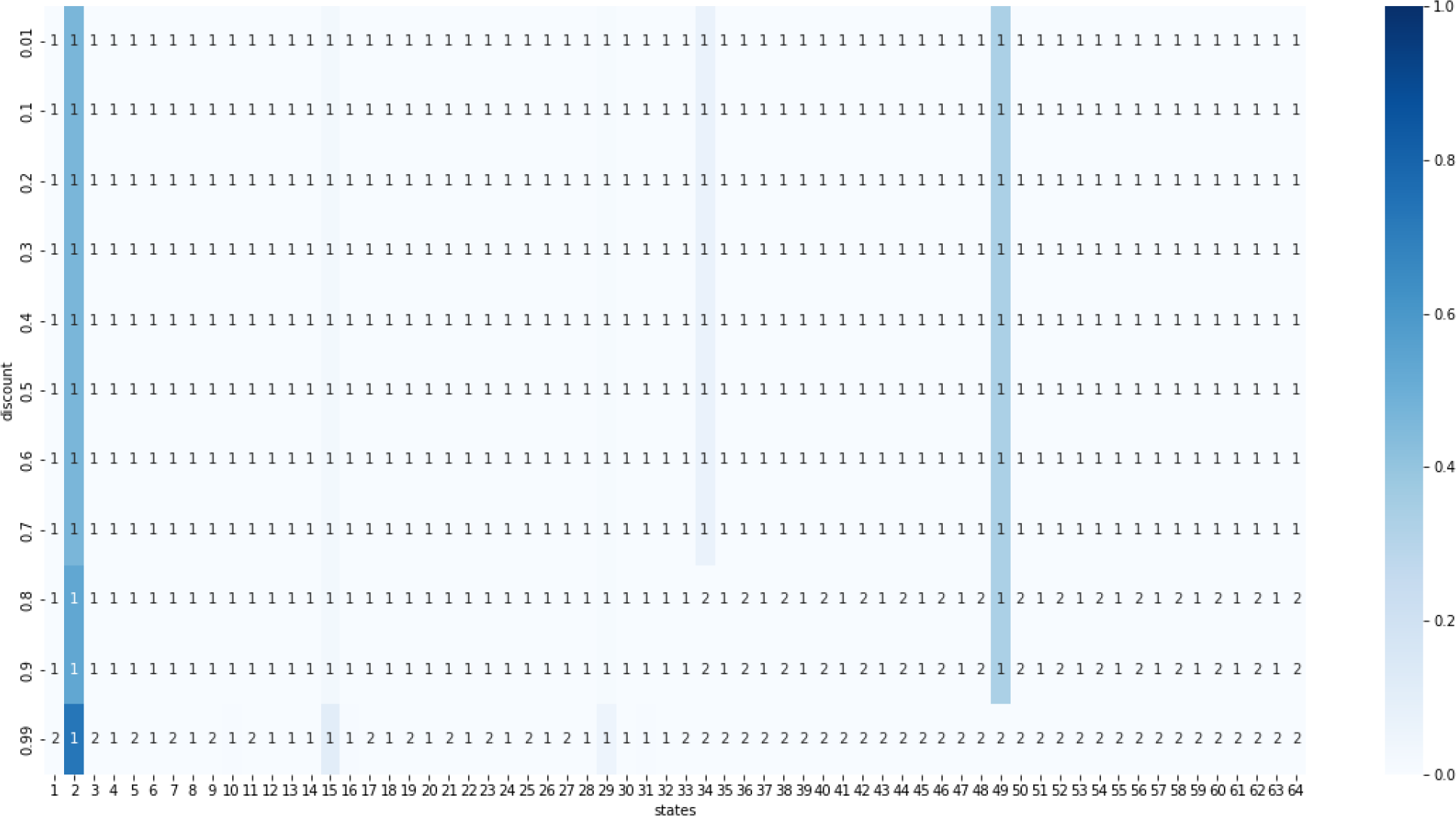
Heatmap of the stationary distribution given propensities from Table 1 using eight-step optimal policies from the value iteration method. The states are on the *x*-axis and the *y*-axis gives the discount used to find the policy. The policies for these choice of parameters are always ineffective.

## References

[1] Alan Veliz-Cuba and Brandilyn Stigler. Boolean models can explain bistability in the lac operon. Journal of computational biology, 18(6):783–794, 2011.

[2] David J Wooten, Jorge Gómez Tejeda Zañudo, David Murrugarra, Austin M Perry, Anna Dongari-Bagtzoglou, Reinhard Laubenbacher, Clarissa J Nobile, and Réka Albert. Mathematical modeling of the candida albicans yeast to hyphal transition reveals novel control strategies. PLoS computational biology, 17(3):e1008690, 2021.

[3] Daniel Plaugher and David Murrugarra. Modeling the pancreatic cancer microenvironment in search of control targets. Bulletin of Mathematical Biology, 83(11):1–26, 2021.

[4] Daniel Plaugher, Boris Aguilar, and David Murrugarra. Uncovering potential interventions for pancreatic cancer patients via mathematical modeling. Journal of theoretical biology, 548:111197, 2022.

[5] Pedro Márquez-Zacarías, Rozenn M Pineau, Marcella Gomez, Alan Veliz-Cuba, David Murrugarra, William C Ratcliff, and Karl J Niklas. Evolution of cellular differentiation: from hypotheses to models. Trends in Ecology & Evolution, 36(1):49–60, 2021.

[6] Tomáš Helikar, Bryan Kowal, Sean McClenathan, Mitchell Bruckner, Thaine Rowley, Alex Madrahimov, Ben Wicks, Manish Shrestha, Kahani Limbu, and Jim A Rogers. The cell collective: toward an open and collaborative approach to systems biology. BMC systems biology, 6(1):1–14, 2012.

[7] Ilya Shmulevich, Edward R. Dougherty, Seungchan Kim, and Wei Zhang. Probabilistic boolean networks: a rule-based uncertainty model for gene regulatory networks. Bioinformatics, 18(2):261–274, 2002.

[8] Ilya Shmulevich and Edward R. Dougherty. Probabilistic Boolean Networks - The Modeling and Control of Gene Regulatory Networks. SIAM, 2010.

[9] O Golinelli and B Derrida. Barrier heights in the kauffman model. Journal De Physique, 50(13):1587–1601, 1989.

[10] Panuwat Trairatphisan, Andrzej Mizera, Jun Pang, Alexandru Adrian Tantar, Jochen Schneider, and Thomas Sauter. Recent development and biomedical applications of probabilistic boolean networks. Cell communication and signaling, 11(1):1–25, 2013.

[11] Dávid Deritei, Nina Kunšič, and Péter Csermely. Probabilistic edge weights fine-tune boolean network dynamics. bioRxiv, 2022.

[12] David Murrugarra, Alan Veliz-Cuba, Boris Aguilar, Seda Arat, and Reinhard Laubenbacher. Modeling stochasticity and variability in gene regulatory networks. EURASIP Journal on Bioinformatics and Systems Biology, 2012(1):5, 2012.

[13] Richard S Sutton and Andrew G Barto. Reinforcement learning: An introduction, volume 1. MIT press Cambridge, 1998.

[14] Dimitri Bertsekas. Reinforcement learning and optimal control. Athena Scientific, 2019.

[15] Mohammadmahdi R Yousefi, Aniruddha Datta, and Edward R Dougherty. Optimal intervention strategies for therapeutic methods with fixed-length duration of drug effectiveness. Signal Processing, IEEE Transactions on, 60(9):4930–4944, 2012.

[16] Boris Aguilar, Pan Fang, Reinhard Laubenbacher, and David Murrugarra. A near-optimal control method for stochastic boolean networks. Letters in Biomathematics, 7(1):67–80, May 2020.

[17] Dimitri P. Bertsekas. Dynamic Programming and Optimal Control. Athena Scientifik, 2005.

[18] David Murrugarra and Boris Aguilar. Modeling the stochastic nature of gene regulation with boolean networks. In Algebraic and Combinatorial Computational Biology, pages 147–173. Elsevier, 2019.

[19] Assieh Saadatpour, István Albert, and Réka Albert. Attractor analysis of asynchronous boolean models of signal transduction networks. J Theor Biol, 266(4):641–56, Oct 2010.

[20] CM Grinstead and JL Snell. Introduction to probability: American mathematical society: Providence. Rhode Island, United States, 2012.

[21] David A Levin and Yuval Peres. Markov chains and mixing times, volume 107. American Mathematical Soc., 2017.

[22] David Murrugarra, Alan Veliz-Cuba, Boris Aguilar, and Reinhard Laubenbacher. Identification of control targets in boolean molecular network models via computational algebra. BMC Syst Biol, 10(1):94, Sep 2016.

[23] Jorge G T Zañudo and Réka Albert. Cell fate reprogramming by control of intracellular network dynamics. PLoS Comput Biol, 11(4):e1004193, Apr 2015.

[24] Jorge Gomez Tejeda Zañudo, Gang Yang, and Réka Albert. Structure-based control of complex networks with nonlinear dynamics. Proceedings of the National Academy of Sciences, 114(28):7234–7239, 2017.

[25] Uri Alon. An Introduction to Systems Biology. Chapman and Hall/CRC, Boca Raton, 2019.

[26] Michael B Elowitz and Stanislas Leibler. A synthetic oscillatory network of transcriptional regulators. Nature, 403(6767):335–338, 2000.

